# PHAROS: turning single-cell perturbation models into target-directed drug-combination screens

**DOI:** 10.64898/2026.09.08.749477

**Authors:** Jon Bezney, Carlo Ruggeri, Federico Borra, Lei S. Qi, Francesca Buffa, Lars M. Steinmetz

## Abstract

Combination therapies are central to cancer treatment, but exhaustive screening is impractical. We introduce PHAROS, a framework that turns a pretrained single-cell perturbation model into a target-directed search engine for drug combinations. PHAROS predicts how a cell population changes under a drug, one drug at a time, then chains these predictions together to simulate drug combinations. It scores each simulated outcome against the desired target state and uses a search algorithm to find the most promising combinations, all without retraining the underlying model. Across two independent combinatorial perturbation datasets, PHAROS recovered exact or mechanism-matched two-drug responses in cell lines, both seen and unseen during model training. Its rankings were specific to the requested conversion and were not explained by single-drug effects, additive effects, or shared mechanism of action. In exploratory analyses of patient-derived metastatic HR+/HER2 breast tumors and basal cell carcinoma (BCC), PHAROS prioritized FDA-approved regimens, distinguished combinations by their predicted tumor-versus-immune objective profiles, and nominated pathway-level hypotheses, while explicitly identifying both tumor cohorts as outside the model’s supported distribution. PHAROS provides a modular route from pretrained virtual-cell models to inverse, single-cell combination-screening platforms.

## 1 Introduction

Predicting the response to a perturbation is only one part of the virtual-cell problem. Therapeutic discovery more often asks the inverse question: which interventions are most likely to move a heterogeneous cellular population from its current state toward a desired state? This question is especially challenging for combination therapy, where the search space expands with each additional drug, dose, order, and cellular context.

Single-cell perturbation models have advanced rapidly, from latent-variable and neural optimal-transport methods to compositional and foundation-model approaches that predict responses across held-out doses, perturbations, and treatment-context combinations ^1–9^. However, most methods are evaluated as forward predictors: a perturbation is specified, and the model predicts the resulting cell state. Methods that model drug combinations at single-cell resolution typically require training, adaptation, or perturbational examples from the experimental system being evaluated ^5,10,11^. Related inverse-design methods identify gene targets from matched disease-treatment data ^12^, but cannot prioritize drug combinations or account for single-cell heterogeneity. Signature-retrieval tools prioritize perturbations from bulk profiles without modeling cell-type-specific responses^13^. Thus, the setting addressed here remains unmet: (1) frozen cross-dataset inference, (2) single-cell prediction of drug combinations, and (3) target-directed selection from source and target state distributions.

Here we introduce PHAROS, an admissibility-controlled framework that converts a pretrained single-cell perturbation model from a forward response predictor into a target-directed combinatorial search engine. Admissibility evaluates the model’s expected reliability for a particular query: whether the source and target states are sufficiently represented within the pretrained model’s learned support and whether their separation can be resolved in the embedding space. It is not a claim about biological feasibility or therapeutic efficacy. A biologically achievable conversion may be inadmissible if the relevant states lie outside the model’s training distribution, whereas an admissible conversion remains a prediction that requires experimental validation. PHAROS embeds source and target populations in the STATE single-cell foundation model latent space and applies a frozen STATE perturbation model trained on Tahoe-100M single-drug perturbations without retraining or fine-tuning on the query dataset^8,14^. By operating in a representation learned from a broad corpus of cell states, rather than training a predictor only within the dataset of interest, we hypothesized that PHAROS could better transfer to previously unseen datasets and cellular contexts while making the limits of that transfer explicit through admissibility assessment.

It operates in two complementary inverse-design modes. In the hypothesis-driven mode, the user specifies a set of candidate combinations, and PHAROS ranks them according to their predicted ability to achieve the requested source-to-target conversion, that is, to shift the starting (source) cell population so that it more closely resembles the desired (target) cell population. In the open search mode, no candidate combination is specified in advance, and PHAROS searches the broader intervention space for ordered drug sequences predicted to produce the desired conversion (Fig. 1a). Across both modes, PHAROS applies drugs one at a time, where each drug’s predicted effect depends on the cell population’s current state, not on some fixed effect computed in isolation, so that later drugs act on the outcome of earlier ones. It then reduces this representation to the dimensions that best distinguish the starting and target populations, and measures how close the resulting predicted population is to the target using an optimal transport distance. To preserve population heterogeneity, PHAROS evaluates each conversion across multiple independently sampled source and target cell batches rather than reducing either population to a single representative profile. The open search mode additionally uses diverse beam search to explore the combinatorial space efficiently (Fig. 1b)^15–17^, whereas the hypothesis-driven mode compares prespecified combinations and mechanism-matched alternatives (drugs sharing the same annotated mechanism of action as a compound of interest, used as a substitute when that exact compound is absent from the model’s vocabulary) against random pairs drawn from the Tahoe vocabulary and separates single-drug, additive two-drug, and treatment-order effects. Before either analysis, PHAROS evaluates task admissibility by estimating an empirical calibration reference, query-state support within the learned perturbation manifold, and source-target separability.

**Figure 1:**
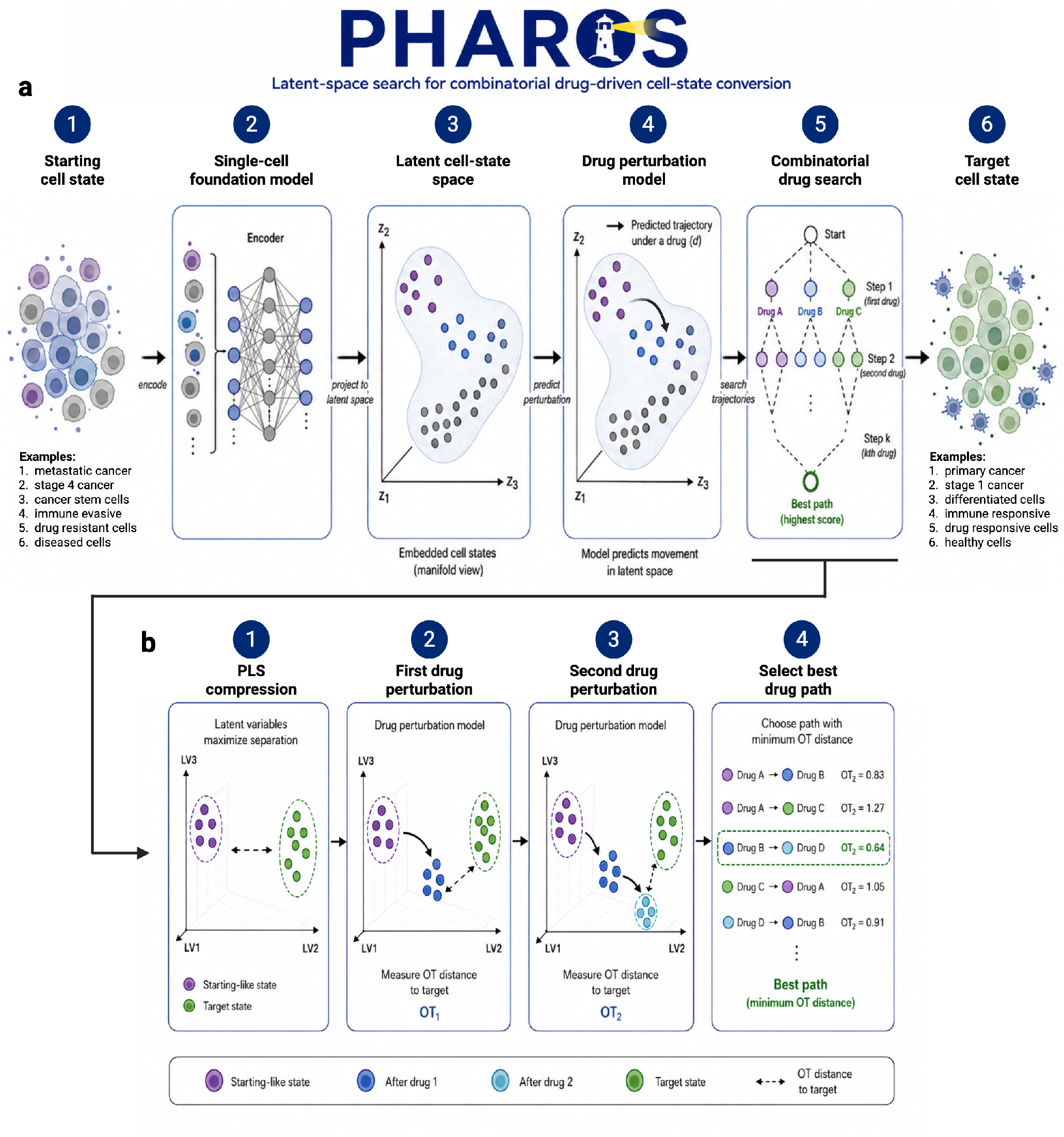
**a,** Schematic showing the workflow of leveraging combinatorial perturbations to convert a starting cell state into a desired target cell state in latent space. A single-cell foundation model is used to embed the cells. A drug perturbation model is used to predict the effect in latent space. A diverse beam search is used to select sequential drug perturbations. **b,** First, starting cells are given a single perturbation and ranked according to distance to target cell. Second, additional perturbations are applied sequentially. The final path is selected according to minimum Sinkhorn OT distance to target.

Because no existing method operates under the same information regime or solves the same target-directed combination-selection task, we evaluated open-search recovery against a beam-matched Monte Carlo null that preserved the size and basic constraints of the search while assigning perturbations independently of the target and predicted scores (Methods). Across two independent combinatorial perturbation datasets, PHAROS ranked experimentally observed or mechanism-matched two-drug responses among the leading candidates, including in cell lines absent from perturbation-model training. In exploratory patient-tumor analyses, PHAROS prioritized FDA-approved metastatic breast-cancer regimens within the top 1% of screened pairs without using approval status during ranking, revealed regimen-specific trade-offs across malignant and lymphoid cell-state objectives, and nominated PLK1- and JAK–STAT-directed hypotheses for basal cell carcinoma (BCC), while explicitly identifying these states as extrapolative. Together, these findings establish PHAROS as an inverse-design layer for pretrained virtual-cell models: rather than predicting the transcriptional response to a treatment selected in advance, it identifies which drug combinations are predicted to best achieve a specified source-to-target cell-state conversion.

## 2 Results

### 2.1 An admissibility framework evaluates unseen cellular contexts

Applying a frozen perturbation model to new data requires deciding whether its predictions are sufficiently supported for the task at hand. PHAROS therefore begins with a three-part admissibility analysis: (1) it measures, using data the model was actually trained on, how much of the gap between a starting and target cell state a predicted intervention can typically close, as a baseline for how much movement toward a target is realistic to expect; (2) it evaluates whether query states remain supported by the reference embedding manifold, that is, it checks whether the new data resembles anything the model has seen during training, by comparing it to the reference set of training cells; and (3) tests whether the proposed source and target populations are separable enough to give the search a clear direction to work toward (Supplementary Fig. S1a). The first component provides a baseline, an empirical reference for score interpretation, whereas the latter two determine whether a specific search should proceed without additional scrutiny, that is, whether it can be trusted or should be treated as a more tentative, hypothesis-generating result.

We first calibrated in-domain performance using Tahoe-100M cell line-drug pairs spanning 3 cell lines and 379 compounds at 5 µM. For each condition, we applied the corresponding perturbation to control cells and compared the predicted state with the measured response. The central 90% of predicted-to-observed Sinkhorn optimal-transport (OT) distances ranged from 303 to 366 (Supplementary Fig. S1b). Relative to the initial control-to-treated distance, predictions closed 31-44% of the source-to-target gap, with a median of 36% (Supplementary Fig. S1c). Thus, even in-domain predictions moved cells reproducibly toward the observed response without reconstructing it exactly. We use these distributions as calibration references rather than absolute success thresholds.

We next applied the support analysis to an external two-drug A549 dataset^5^. Untreated cells fell within the core Tahoe-100M reference distribution, whereas several combination-treated states were classified as boundary conditions (Supplementary Fig. S1d,e). No evaluated state met the criteria for an out-of-distribution call, and nearest reference neighbors were enriched for lung-derived cell lines (Supplementary Fig. S1f), consistent with the A549 origin of the query data. Because A549 is present in Tahoe-100M, this analysis tests transfer across datasets and into previously unseen combination-induced states, rather than transfer to a completely unseen cell identity.

Finally, PHAROS tested whether each source-target pair defined a distinguishable objective. The DMSO and panobinostat-crizotinib populations formed separable neighborhoods and passed the predefined purity criterion (Supplementary Fig. S1g). By contrast, the DMSO and givinostat-crizotinib populations were not reliably separated (Supplementary Fig. S1h). Failure of this gate does not imply absence of a biological effect; it indicates that the current representation does not resolve the requested conversion well enough for confident target-directed search. The admissibility framework therefore separates supported conversions from boundary and poorly resolved cases that require additional scrutiny.

### 2.2 PHAROS recovers exact and mechanism-matched combinations in an external A549 dataset

We evaluated PHAROS on three positive-control conversions from the external A549 combination dataset of Lotfollahi et al ^5^. In the first case, both measured compounds, panobinostat and crizotinib, were present in the perturbation-model vocabulary, enabling exact recovery. In the other two cases, panobinostat was represented but the second measured compound was not, so we tested mechanism-matched proxies: resveratrol, a SIRT1 activator, for SRT-3025, and pimitespib, an HSP90 inhibitor, for alvespimycin (Fig. 2a). These cases distinguish exact compound recovery from mechanism-level transfer to compounds absent from the model vocabulary.

**Figure 2:**
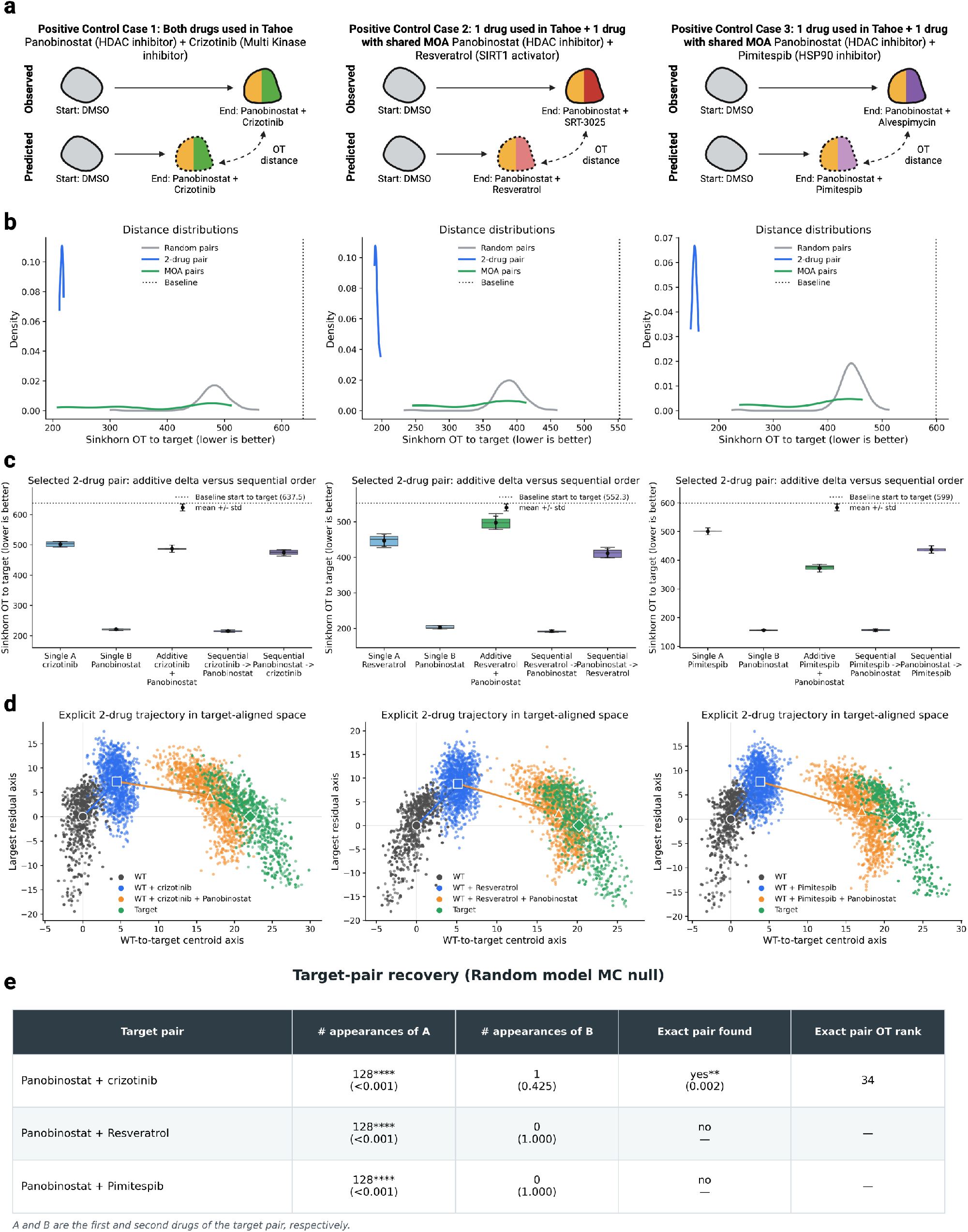
**a,** Schematic of observed versus predicted starting cell to target cell conversions in unseen A549 dataset. Left: panobinostat and crizotinib. Middle: panobinostat and SIRT1 activator. Right: panobinostat and HSP90 inhibitor. **b,** Distribution of Sinkhorn OT distance to target under 2-drug predicted perturbations comparing 100 random pairs (grey), the exact 2-drug pair (blue), and all pairs with the same MOA (green). For each unordered drug pair, we evaluated all tested concentration combinations and both treatment orders, and retained the configuration with the lowest OT distance to the target. All 2-drugs were ran across 5 batches of starting and target cells. Dotted line is baseline OT distance between starting cell state and target cell state. **c,** Boxplot of OT distance to target of starting cell state under multiple perturbation conditions of the known 2-drug combination including: single drug A, single drug B, additive displacement vectors of individual drug A and B, sequential A then B, sequential B then A. **d,** Projected sequential trajectory onto target-aligned plane. The horizontal axis connects the start and target population centroids and therefore represents movement along the desired conversion direction. The vertical axis captures the largest remaining source of variation after removing this start-to-target direction, revealing deviations from a direct trajectory. Observed DMSO WT (black), predicted WT + drug 1 (blue), predicted WT + drug 1 + drug 2 (orange), observed target (green). **e,** Target-pair recovery table for the three drug combinations. # appearances of A indicates the number of recovered paths containing the first drug in the target pair, while # appearances of B indicates the corresponding number for the second drug. Two-drug paths were obtained from the respective open search with a search width of 128. Exact pair found indicates whether both drugs in the target pair occurred together in at least one recovered path. When the exact pair was recovered, Exact-pair OT rank reports its rank according to optimal transport distance to the target, with lower numerical ranks correspond to lower OT distances (rank 1 is best). Monte Carlo (p)-values under a random null model are reported in parentheses.

In the hypothesis-driven mode, for each conversion, DMSO-treated cells defined the source and the measured two-drug population defined the target. The exact or mechanism-matched pair outperformed at least 99% of 100 sampled random controls in all three cases and closed 66%, 65%, and 74% of the initial source-to-target OT distance, respectively (Fig. 2b). Because the hypothesis-driven mode evaluates a specified pair against the broader perturbation background, it also provides an empirical estimate of whether that pair is sufficiently enriched to be recoverable by the open search. Here, the observed percentile ranks placed each positive-control pair near the top of the candidate space, indicating that the corresponding open search, with a sufficiently high width parameter, should be capable of prioritizing it. All three conversions exceeded the central 90% of the Tahoe single-drug calibration despite requiring sequential two-drug prediction. The same pairs did not retain this advantage when the conversion direction was reversed, indicating that recovery was specific to the requested source-to-target task (Supplementary Fig. S2). Mechanism-matched candidate pairs were also shifted toward lower OT distances than random controls, but with substantial variability across candidates (Fig. 2b).

Panobinostat alone accounted for much of the target-directed displacement, consistent with the measured data in which panobinostat-treated cells overlapped strongly with the corresponding two-drug populations (Supplementary Fig. S3). Thus, these controls are biologically dominated by panobinostat, but PHAROS still required state-dependent sequential composition to reproduce the best two-drug predictions. In all three cases, the sum of the two single-drug displacement vectors did not outperform the better of the two single-drug effects (Fig. 2c). In these specific conversions, intervention order at inference mattered. Sequences in which panobinostat was applied second yielded a lower OT distance to the experimental target generated by co-administration of the two drugs.

Target-aligned trajectory analysis clarified how the sequential predictions moved through the embedding space. The first transition generally produced modest progress along the start-to-target axis and, in some cases, a residual off-axis displacement. The second transition produced most of the target-directed movement and reduced the residual displacement introduced by the first step (Fig. 2d).

We next evaluated PHAROS open search capabilities to find promising drug pairs that move the source A549 control-like population toward each of the three positive-control populations generated by two-drug perturbations. The search considered 379 drugs at three concentrations, yielding 1,137 drug-concentration labels and 1,289,358 possible ordered two-step trajectories after prohibiting reuse of the same base drug within a path (Fig. 3e).

**Figure 3:**
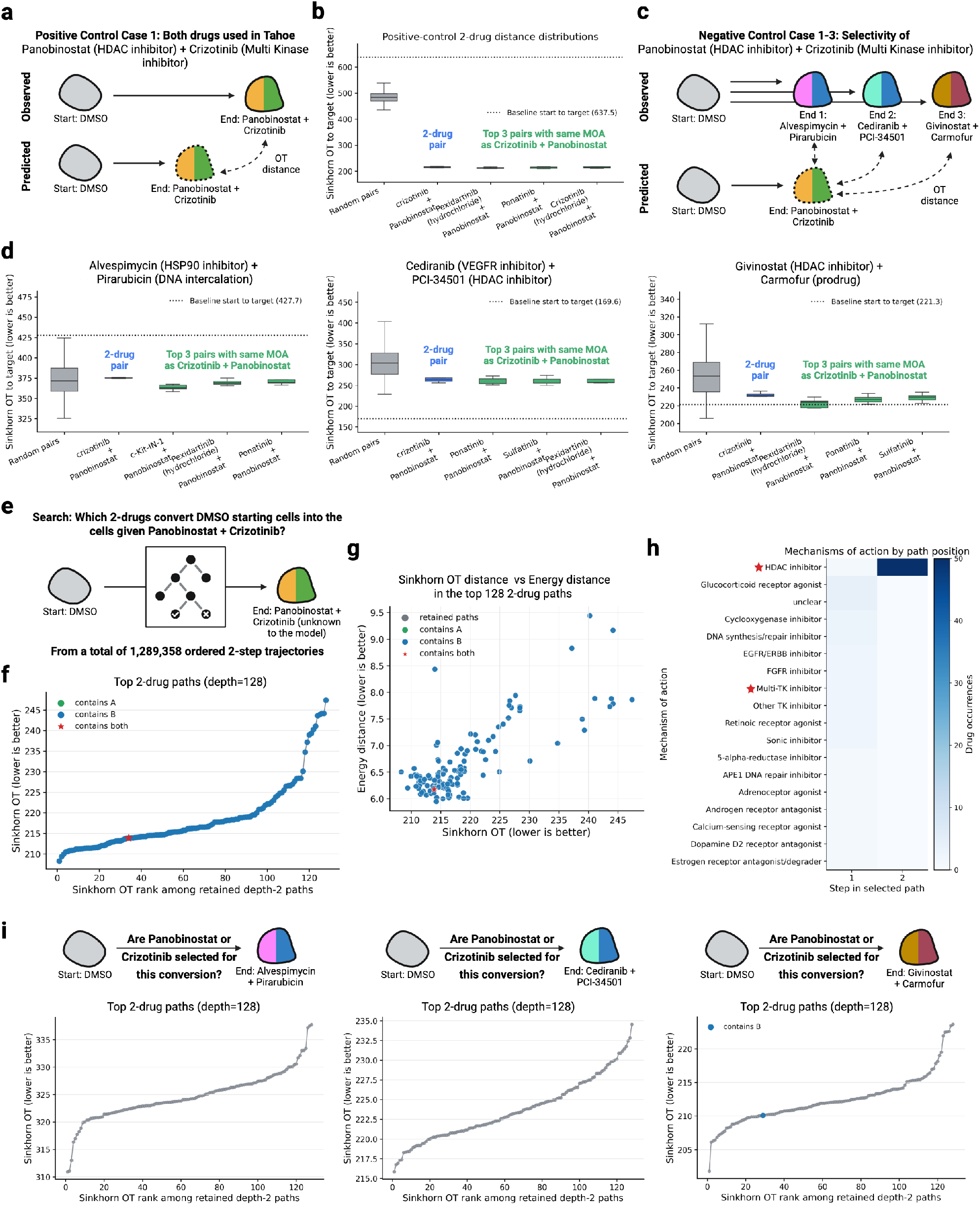
**a,** Schematic of observed versus predicted where both drugs, panobinostat + crizotinib, are seen. **b,** Observed matches predicted. Distributions of OT distance to target comparing 100 random 2-drugs (grey), exact panobinostat + crizotinib (blue), and top 3 pairs with matching MOA as crizotinib + panobinostat (green). For each unordered drug pair, we evaluated all tested concentration combinations and both treatment orders, and retained the configuration with the lowest OT distance to the target. All 2-drugs were ran across 5 batches of starting and target cells. Dotted line indicates baseline start-to-target distance. **c,** Schematic demonstrating 3 counterfactual examples where the predicted pair, panobinostat + crizotinib, is compared against the observed pairs of alvespimycin + pirarubicin, cediranib + PCI34501, and givinostat + carmofur. **d,** 3 scenarios where observed does not match predicted. Distributions of OT distance to target comparing 100 random 2-drugs (grey), exact panobinostat + crizotinib (blue), and top 3 pairs with matching MOA as crizotinib + panobinostat (green). For each unordered drug pair, we evaluated all tested concentration combinations and both treatment orders, and retained the configuration with the lowest OT distance to the target. All 2-drugs were ran across 5 batches of starting and target cells. Dotted line indicates baseline start-to-target distance. **e,** Schematic of beam search over all potential 2-drug pairs to select the top paths to convert the observed starting DMSO cell state into the observed target cell state (panobinostat + crizotinib). **f,** Line plot of top 128 ranked 2-drug paths (x-axis) versus OT distance to target (y-axis). Green indicates crizotinib present in path, Blue indicates panobinostat present in path, red star indicates both present. **g,** Scatterplot comparing the energy distance to the target with the Sinkhorn OT distance to the target across the 128 two-drug paths recovered by the open search. **h,** Heatmap of MOA frequencies across each step of the 2-step trajectory for the top 50 paths. Red star denotes MOAs of panobinostat and crizotinib. **i,** Line plot of top 128 ranked 2-drug paths (x-axis) versus OT distance to target (y-axis) for the 3 counterfactual examples. Green indicates crizotinib present in path, blue indicates panobinostat present in path, red star indicates both present.

With depth two and beam width 128, PHAROS recovered panobinostat-crizotinib at rank 34 among the retained depth-two paths (Fig. 3f,g). Under the null search, both recovery of the exact drug pair and enrichment of panobinostat beyond that observed in the actual search were highly unlikely (p = 0.002 and p < 0.001, respectively; Fig. 2e). Panobinostat appeared in every retained path, whereas partner drugs were more diverse (Fig. 3f,g). At the mechanism level, the search consistently placed an HDAC inhibitor, usually panobinostat, at the second step and selected a broader range of mechanisms at the first step; multi-tyrosine-kinase inhibitors, including the class containing crizotinib, were the eighth most frequent first-step mechanism (Fig. 3h). Cases 2 and 3 also recovered panobinostat broadly, but not the designated mechanism-matched second compounds (Fig. 2e). This lower partner-level recovery likely reflects two limitations: compounds sharing an annotated mechanism can induce distinct transcriptional responses, and beam search can prune an enabling first perturbation if its benefit emerges only after the second step.

### 2.3 PHAROS prioritization is specific to the requested cell-state conversion

An inverse-design method should not simply identify drug pairs that move cells broadly across latent space. It should prioritize a pair when its measured response defines the target and lose that advantage against unrelated targets. We tested this requirement using panobinostat-crizotinib, the positive-control pair for which both compounds were present in the model vocabulary (Fig. 3a).

When the measured panobinostat-crizotinib population was used as the target, the exact pair and the three highest-ranked mechanism-matched pairs fell in the extreme lower tail of the random-control distribution and reduced OT distance relative to the untreated baseline (Fig. 3b). We then scored the same candidate pairs against three mismatched two-drug target states generated by alvespimycin-pirarubicin, cediranib-PCI34501, and givinostat-carmofur (Fig. 3c). Under these counterfactual targets, panobinostat-crizotinib lost its enrichment and struggled to outperform random drug pairs in OT distance to the target (Fig. 3d). The other positive-control cases showed the same pattern (Supplementary Fig. S4), indicating that PHAROS rankings were conditional on the requested conversion.

Residual cross-target activity was biologically structured rather than random. Several high-scoring candidates shared the HDAC-inhibitor mechanism of panobinostat, especially when the mismatched target contained givinostat or PCI34501, another HDAC-inhibitor (Fig. 3d). These effects were stronger in the conversion-focused compressed space than in the full embedding space, consistent with compression emphasizing dimensions that separate the requested source and target (Supplementary Fig. S5). We therefore interpret these controls as revealing a useful trade-off: conversion-focused scoring increases sensitivity to target-aligned movement but requires mismatched-target controls to guard against over-interpreting mechanistically related states as exact recovery.

Finally, PHAROS specificity carried forward also within the open search framework. Neither drug showed meaningful enrichment among retained paths for the mismatched targets: crizotinib was entirely absent, and panobinostat appeared in only one path across the three searches (Fig. 3i).

### 2.4 PHAROS recovers combinations in cell states absent from perturbation-model training

We next tested whether PHAROS could recover drug combinations in cellular contexts absent from perturbation-model training. We applied the frozen framework to the glioblastoma combination dataset of McFaline-Figueroa et al. ^18^, which includes A172 cells represented in Tahoe-100M and U87MG and T98G cells absent from training. For each cell line, we evaluated five combinations in which both compounds were present in the model vocabulary and five in which one compound required a mechanism-matched proxy. Across the three cell lines, these two panels comprised 30 distinct two-drug benchmarking conversions.

The admissibility analysis separated reference support from conversion resolvability and identified this dataset as a stringent stress test for PHAROS. All three cell lines and their perturbation states were classified as core members of the Tahoe-100M embedding reference and localized near brain-derived cancer cell lines (Supplementary Fig. S6). Despite this reference support, every source-target conversion failed the neighborhood-separability criterion because untreated and treated populations were poorly resolved (Supplementary Fig. S7). This limited resolution coincided with a median of only approximately 477-717 detected genes per cell, suggesting that low transcript recovery obscured perturbation-induced state differences. In addition, some drug-concentration conditions contained fewer than 10 cells, requiring us to pool multiple concentrations to define target populations of sufficient size (Supplementary Fig. S8). We therefore applied a high-sensitivity analysis that selects diverse source and target batches with stronger baseline separation (Fig. 4a; Methods). Because this procedure conditions on separability, the resulting scores quantify performance within informative subpopulations rather than across the complete cell populations. Thus, rather than representing a favorable benchmark, this dataset deliberately tests the limits of PHAROS under sparse cell sampling, low gene-detection sensitivity, poorly resolved treatment states, and cellular contexts absent from perturbation-model training.

**Figure 4:**
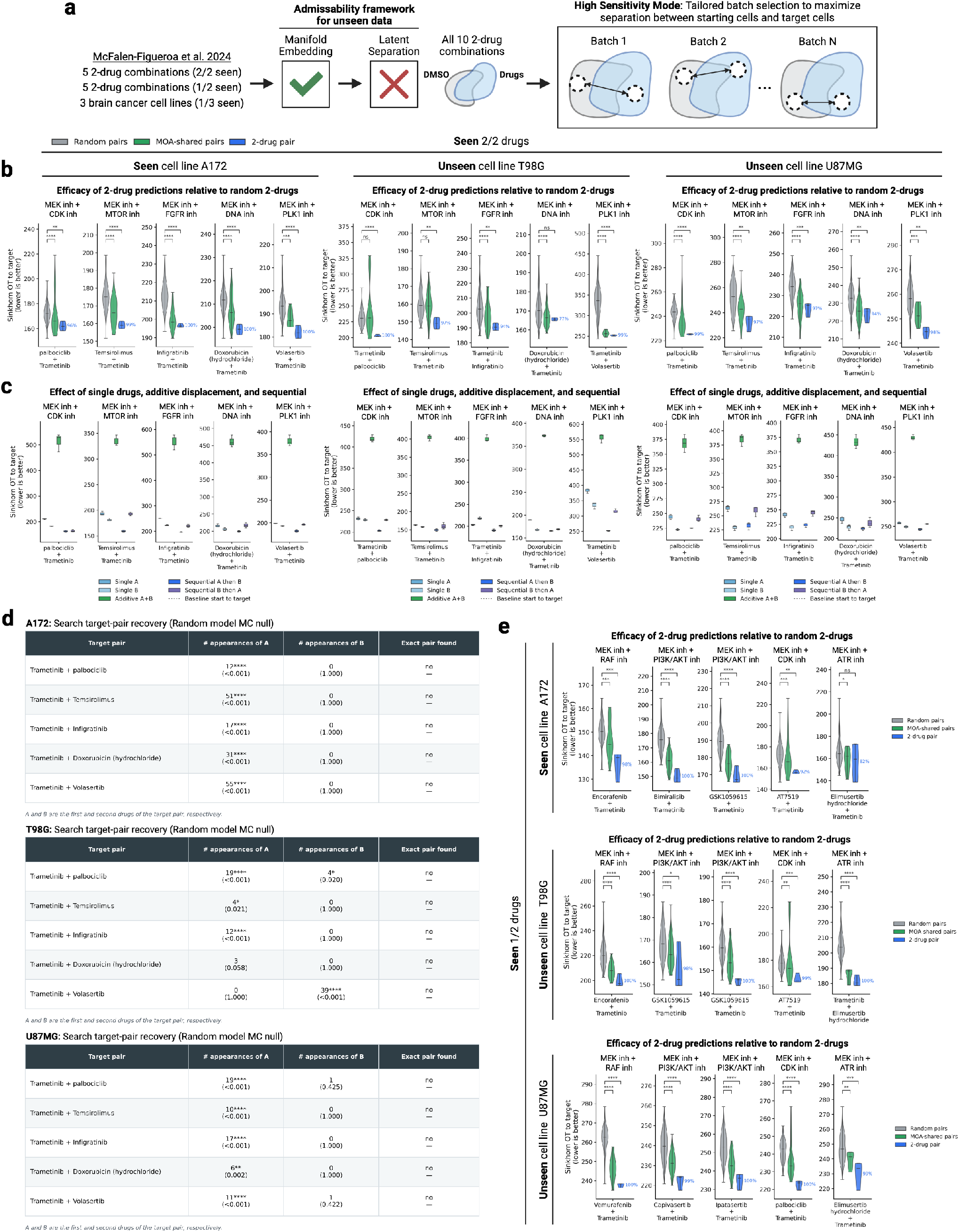
**a,** Schematic of PHAROS application to unseen 2-drug perturbation data across 3 brain cancer cell lines. Explanation of high sensitivity mode to select batches of cells to maximize separation of start and target cell states. **b,** Distributions of OT distances to the target for 100 random two-drug control pairs (grey), all drug pairs with matching MOAs (green), and the exact target pair (blue), across five combinations for which both drugs were present in Tahoe-100M. For each unordered drug pair, we evaluated all tested concentration combinations and both treatment orders, and retained the configuration with the lowest OT distance to the target. All 2-drugs were ran across 3 batches of starting and target cells. Results are shown for three cell lines: A172 (seen), T98G (unseen), and U87MG (unseen). Statistical significance was assessed using one-sided Mann–Whitney (U) tests with Benjamini–Hochberg correction for multiple comparisons. **c,** Distributions of OT distances to the target comparing single-drug perturbations, additive displacement predictions, and sequentially ordered perturbations across five drug combinations for which both drugs were seen. Results are shown for three cell lines: A172 (seen), T98G (unseen), and U87MG (unseen). **d,** Target-pair recovery tables for the five combinations for which both drugs were present in Tahoe-100M. # appearances of A indicates the number of recovered paths containing the first drug in the target pair, while # appearances of B indicates the corresponding number for the second drug. Two-drug paths were obtained from the respective open search with a search width of 128. Exact pair found indicates whether both drugs in the target pair occurred together in at least one recovered path. When the exact pair was recovered, exact-pair OT rank reports its rank according to optimal transport distance to the target, with lower numerical ranks correspond to lower OT distances (rank 1 is best). Monte Carlo (p)-values under a random null model are reported in parentheses. Results are reported for the three cell lines: A172 (seen), T98G (unseen), and U87MG (unseen). **e,** Distributions of OT distances to the target for 100 random two-drug control pairs (grey), all drug pairs with matching MOAs (green), and best pair with matching MOA (blue), across five combinations for which one out of two drugs was present in Tahoe-100M. For each unordered drug pair, we evaluated all tested concentration combinations and both treatment orders, and retained the configuration with the lowest OT distance to the target. All 2-drugs were ran across 3 batches of starting and target cells. Results are shown for three cell lines: A172 (seen), T98G (unseen), and U87MG (unseen). Statistical significance was assessed using one-sided Mann–Whitney (U) tests with Benjamini–Hochberg correction for multiple comparisons.

In A172, the cell line represented during perturbation-model training, the five exact-vocabulary combinations outperformed 96-100% of sampled ordered random controls, where higher percentages indicate more random controls with equal or greater, and therefore worse, OT distance (Fig. 4b). Mechanism-transfer cases were less consistent, ranging from 82% to 100%, with performance reported using, for each two-drug pair, the best-performing mechanism-matched proxy as determined by final OT distance (Fig. 4e). Unlike the A549 positive controls, these predictions were not reproduced by either constituent drug alone or by additive single-drug displacement vectors (Fig. 4c). In the open searches with beam width 128, PHAROS did not recover any of the five exact 2-drug pairs (Fig. 4d). However, in the hypothesis-driven mode, most of these pairs ranked near the 99th percentile among random controls, suggesting that the absence of exact recovery reflected limited beam coverage rather than low conversion scores. Extrapolating from the search space size 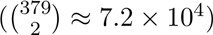 unordered drug pairs, concentrations excluded), increasing the beam width to roughly 1% of that space (720 paths) would be expected to retain most pairs with hypothesis-driven modality performance at or above the 99th percentile. Still, in all open searches trametinib appeared in the retained beam more often than expected under the random-beam null (p < 0.001 for # of appearances of A in all searches; Fig. 4d).

PHAROS also prioritized combinations in T98G, a cell line absent from training. Three out of five exact-vocabulary combinations outperformed 97% of random controls, whereas the remaining outperformed 94% and 77%. Mechanism-transfer cases outperformed 98-100% of random controls (Fig. 4e). Again, neither single-drug predictions nor additive vectors reproduced the sequential two-drug effects (Fig. 4c). Similarly to A172, the open searches did not recover any of the five exact two-drug pair and generally showed significant enrichment of one of the two drugs relative to the random-beam null. For the trametinib–palbociclib target pair, the retained top paths were significantly enriched for both drugs (p 0.02 for each), indicating that PHAROS recovered perturbational signals associated with both components of the target combination (Fig. 4d).

Results in U87MG, the second unseen cell line, remained within the range observed for T98G. Exact-vocabulary combinations outperformed 94-99% of random controls, and mechanism-transfer combinations outperformed 99-100% (Fig. 4b,e). For three of the five exact two-drug target pairs, the best-performing single drug achieved better performance than the best-performing sequential combination (Fig. 4c). In general, these results extend PHAROS from cross-dataset transfer to cell-state transfer and demonstrate its ability to recover substantial biological signal despite limitations in dataset quality. Predictions remained target- and context-dependent even when the cell line was absent from model training.

### 2.5 Exploratory prioritization of approved combinations in metastatic HR+/HER2 breast cancer

Having evaluated PHAROS in external cell-line perturbation datasets, we next asked whether it could generate clinically grounded, patient-specific hypotheses from tumor-derived cell states. We analyzed malignant cells from the breast-cancer atlas of Ozmen et al.^19^, selecting six metastatic HR+/HER2 samples spanning biopsy sites, tumor phenotypes, and prior treatment histories (Fig. 5a,b).

**Figure 5:**
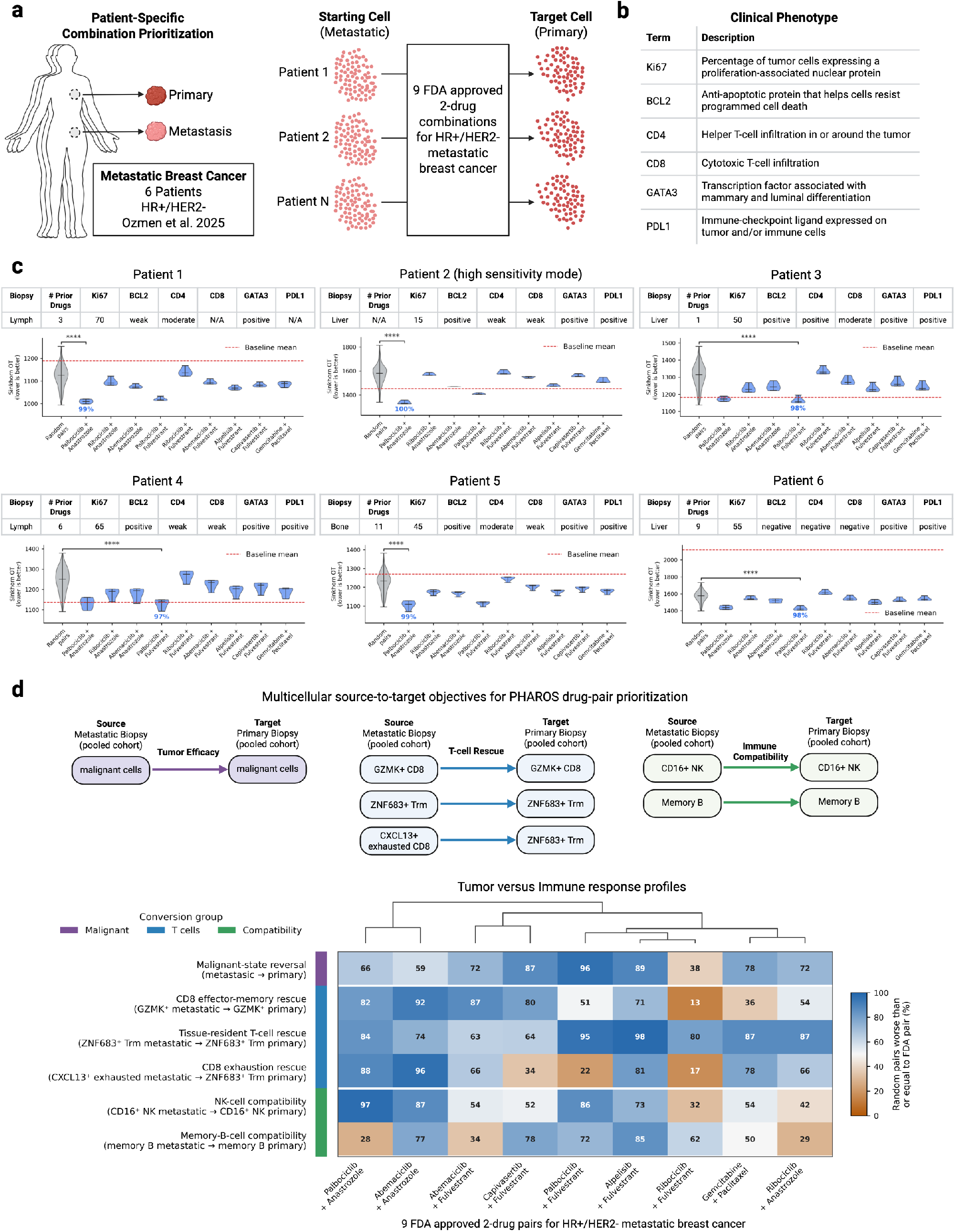
**a,** Schematic of conversion from metastatic malignant-cell populations to matched primary-like malignant-cell reference populations across six metastatic HR+/HER2− breast-cancer samples. **b,** Description of biological markers used to clinically characterize the tumors. **c,** Top: table of clinical phenotype scores. Bottom: distributions of OT distances from start to target comparing 100 random 2-drugs (grey) versus nine FDA-approved two-drug combinations for HR+/HER2− breast cancer (blue). For each unordered drug pair, we evaluated all tested concentration combinations and both treatment orders, and retained the configuration with the lowest OT distance to the target. All 2-drugs were ran across 5 batches of starting and target cells. Dotted line represents baseline start-to-target distance. Significance was assessed using a one-sided Mann-Whitney U test with Benjamini-Hochberg correction on Sinkhorn OT distances, testing whether the control group had greater OT distance. **d,** 9 FDA-approved drug-pair performance across 6 source-to-target objectives. Rows represent malignant-state reversal, three CD8 T-cell rescue objectives, and NK-cell and memory-B-cell compatibility objectives. Each cell shows the percentage of 100 matched random drug pairs with mean Sinkhorn optimal-transport distance worse than or equal to the FDA pair; higher values (blue) indicate more favorable performance relative to random pairs, whereas lower values (orange) indicate poorer relative performance. Drug pairs were hierarchically clustered by their six-conversion response profiles.

For each metastatic sample, the malignant-cell population defined the source, and the most transcriptionally similar primary malignant population in the atlas defined the target. Because the primary and metastatic samples were obtained from different individuals, this target does not represent a matched antecedent tumor. Instead, it provides a standardized proof-of-concept task: identify interventions predicted to shift each metastatic population toward a primary-like malignant state. We reasoned that, because primary breast cancers have an overall survival exceeding 90%, whereas metastatic breast cancer is substantially more difficult to treat and has an overall survival of approximately 25%, a shift toward a primary-like state could serve as an exploratory proxy for a potentially more favorable tumor phenotype ^20,21^. This analysis therefore tests whether PHAROS can generate hypotheses for drug combinations that may redirect difficult-to-treat metastatic cell states toward states associated with earlier-stage disease.

The admissibility analysis classified most primary and metastatic populations as out of distribution relative to the Tahoe-100M reference, consistent with the biological distance between patient-derived tumors and cancer cell lines (Supplementary Fig. S9). Nevertheless, nearest reference neighbors were enriched for breast-cancer cell lines, and five of six source-target pairs were separable in the embedding space (Supplementary Fig. S10). Patient two required the high-sensitivity analysis introduced above. We therefore treated this cohort as an exploratory extrapolation test rather than a validated prediction setting. Importantly, the admissibility framework makes this evidentiary boundary explicit, preventing predictions in patient-derived samples from being interpreted with the same confidence as analyses supported by the cell-line training distribution.

Within the model vocabulary, nine combinations were FDA-approved for HR+/HER2 metastatic breast-cancer settings. Approval status defined the clinically constrained evaluation panel but was not used during ranking. We hypothesized that combinations with established activity in metastatic disease might also produce perturbational shifts aligned with transcriptional differences between metastatic and primary-like tumor states. In the hypothesis-driven modality, the best-scoring approved combination outperformed 97-100% of random controls across the six samples, with a median of 99% (Fig. 5c). Rankings varied across patients, with two distinct regimens selected as the leading combination, illustrating the potential for patient-state-specific prioritization. These rankings were not reproduced by additive perturbation vectors (Supplementary Fig. S11).

We next asked whether an open search over the broader drug-combination space could identify additional biologically informative hypotheses for the same metastatic-to-primary-like conversion task. Out of 139 total MOAs, PHAROS converged on two-drug combinations enriched for DNA synthesis and repair inhibitors together with FGFR and EGFR inhibitors across all 6 patients (Supplementary Fig. S12). FGFR and EGFR signaling have been implicated in breast cancer proliferation, invasion, and chemoresistance, including during metastatic outgrowth, while metastatic cells often exhibit heightened dependence on DNA damage response pathways to tolerate replication stress^22–28^. Consistent with this, DNA repair inhibition is an established therapeutic strategy in breast cancer, particularly in genomically unstable or DDR-deficient tumors^29,30^. Thus, the recurrent prioritization of FGFR/EGFR and DNA repair–directed combinations shows that PHAROS can generate candidate drug pairs that align with known signaling and genome-maintenance dependencies associated with the primary–to–metastatic transition in breast cancer. These results remain hypothesis-generating, but they illustrate how the framework can be used not only to rank existing treatment options but also to propose new combinations for specific cell-state conversion objectives.

To assess whether malignant-state reversal coincided with favorable immune-state compatibility, we used PHAROS to define one tumor-intrinsic and five lymphoid source-to-target cell-state objectives. The lymphoid objectives captured restoration of GZMK+ effector-memory and ZNF683+ tissue-resident-memory CD8+ programs, an exhausted-to-resident-memory CD8+ transition, and preservation of CD16+ NK-cell and memory-B-cell states. Alpelisib plus fulvestrant showed the most consistently favorable profile, ranking above most matched random pairs for malignant-state reversal, all three CD8+ T-cell objectives, and both NK- and B-cell compatibility endpoints. In contrast, palbociclib plus fulvestrant produced the strongest malignant-state reversal but limited exhausted-CD8-to-resident-memory rescue, consistent with a tumor-dominant rather than broadly immune-supportive profile. Abemaciclib plus anastrozole showed the converse pattern, with strong lymphoid rankings but more modest malignant-state reversal (Fig. 5d). More broadly, these results show that, within the modeled objectives, FDA-approved combinations can exhibit distinct tumor-versus-immune response profiles. PHAROS enables users to specify biologically precise cell-state conversions and evaluate every candidate combination against a common panel of user-defined objectives. This produces multi-objective, cross-cell-type prioritization profiles that capture predicted alignment with malignant and immune-state objectives and reveal potential cell-type-specific trade-offs.

Together, these results establish a proof-of-concept use case for PHAROS: translating patient-derived tumor states into state-dependent and patient-specific combination hypotheses when perturbation measurements from the relevant patient context are unavailable or impractical to collect. The enrichment of approved combinations within the clinically constrained panel provides retrospective clinical concordance, but it does not demonstrate that the selected treatments would have benefited the corresponding patients. Similarly, the open-search results identify biologically plausible candidate combinations rather than validated therapeutic interventions. The out-of-distribution classification defines this evidentiary boundary and highlights the settings in which experimental validation is most important. Prospective testing in patient-derived models, broader tumor perturbation training data, and matched longitudinal samples will be required to determine whether these rankings predict treatment response or support the development of patient-specific second-line therapies.

### 2.6 Exploratory conversion of immunotherapy-refractory basal cell carcinoma states

We next asked whether PHAROS could identify pharmacologic routes that may prime immunotherapy-refractory malignant cells for PD-1 blockade. We analyzed site-matched single-cell RNA-seq profiles from primary basal cell carcinoma (BCC) biopsies collected before and after anti-PD-1 therapy in 11 patients (6 responders and 5 non-responders; Fig. 6a) ^31^. Malignant cells from responders and non-responders occupied separable post-treatment states in the embedding space (Supplementary Fig. S13). We therefore used post-treatment non-responder malignant cells as the source and post-treatment responder malignant cells as the target, asking which perturbations would shift resistant tumor cells toward the responder-associated malignant-cell state (Fig. 6b).

**Figure 6:**
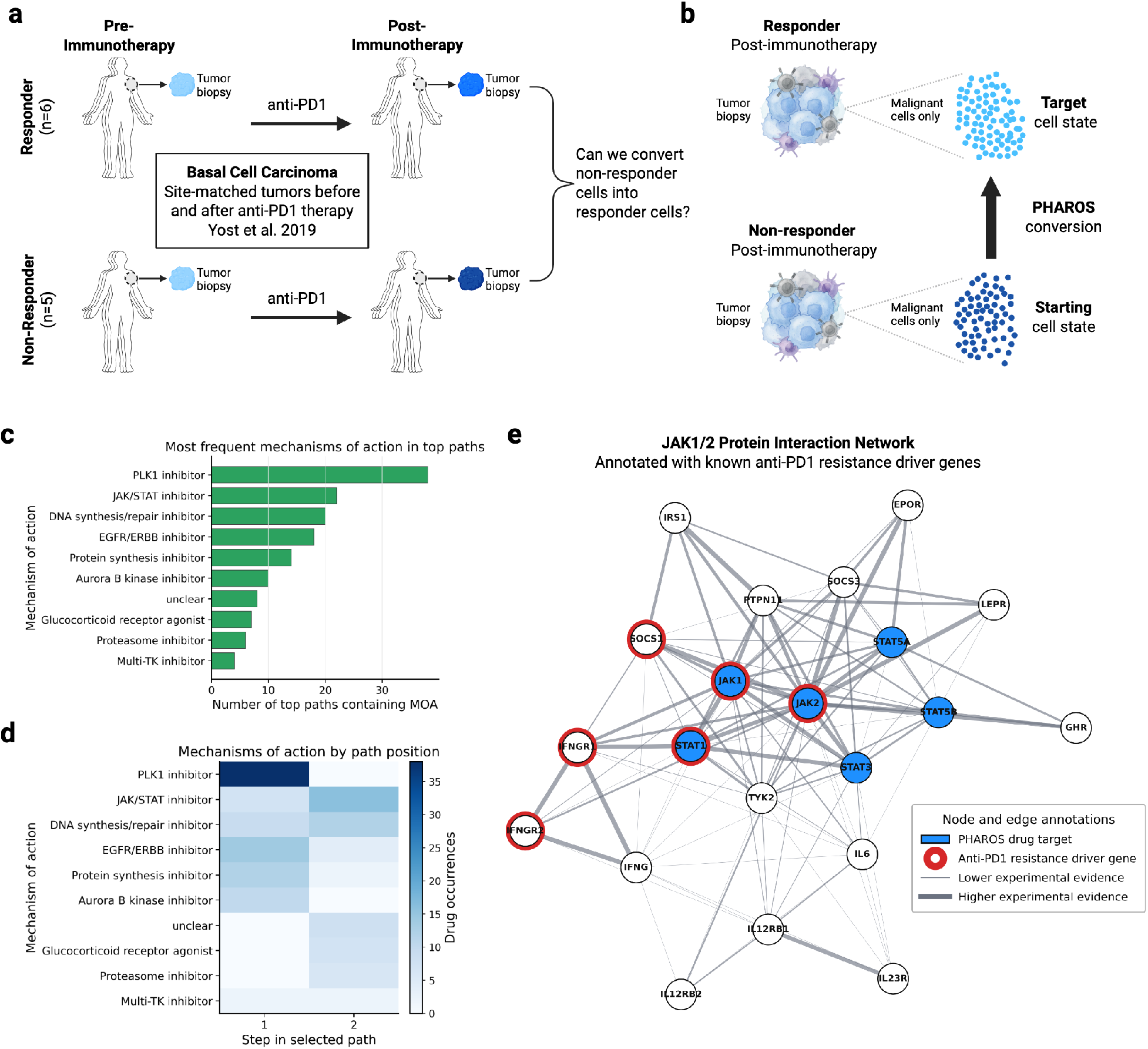
**a,** Study design for site-matched BCC biopsies collected before and after anti-PD-1 therapy from responders (n=6) and non-responders (n=5). **b,** PHAROS conversion objective, from post-treatment malignant cells of non-responders to those of responders. **c,** Mechanism-of-action frequencies derived from the top 100 2-drug trajectories selected by PHAROS in the open-search mode. **d,** Heatmap of mechanism-of-action occurrences per step in the top 100 2-step trajectories selected by PHAROS in the open-search mode. **e,** JAK1/2 protein–protein interaction network from the STRING database. Edge weights denote experimental evidence. Red node outline denotes known driver genes responsible for anti-PD-1 resistance. Blue coloring denotes drug targets from PHAROS selected JAK/STAT inhibitors.

This target is biologically meaningful even though anti-PD-1 acts principally on the immune compartment. Checkpoint blockade requires not only reinvigorated T cells, but also tumor cells that remain visible and susceptible to immune attack^32^. The responder-associated malignant-cell state therefore provides a tumor-cell-intrinsic readout of a context permissive for immune control. PHAROS does not model the immune microenvironment or predict the effect of anti-PD-1 alone, but instead identifies candidate co-treatments that could move non-responder tumor cells toward a state in which immune-mediated killing might be more effective.

In open-search mode, PHAROS recurrently selected PLK1-family and JAK–STAT-pathway inhibitors as the highest-scoring candidates (Fig. 6c,d). The IFN-γ–JAK–STAT axis is among the most strongly supported tumor-cell determinants of checkpoint response across cancers. IFN-γ signaling through IFNGR1/2, JAK1/2, STAT1, and IRF1 induces antigen-processing machinery, MHC class I expression, and adaptive PD-L1 expression. Persistent or dysregulated IFN–JAK–STAT signaling can promote adaptive resistance through inhibitory programs^32–37^. The recurrence of JAK–STAT-targeting compounds therefore identifies this response-defining axis as a plausible route for tumor-cell reprogramming.

The directionality of this prediction is important. Because both loss of productive IFN-γ signaling and chronic, maladaptive pathway activation can impair checkpoint response ^32,36,38–40^. A pathway-level hit does not imply that indiscriminate JAK inhibition will sensitize BCC to anti-PD-1. Rather, PHAROS nominates state-dependent modulation of this axis as a tumor-cell-priming hypothesis whose direction, dose, and schedule must be resolved experimentally in combination with PD-1 blockade.

PLK1 provides a complementary rationale. As a central mitotic kinase, PLK1 regulates mitotic progression and cell-cycle control, and its high expression is broadly associated with proliferative and therapy-resistant tumor programs ^41–44^. Beyond direct cytotoxicity, PLK1 inhibition can induce immunogenic cell death, promote dendritic-cell activation and antigen presentation, and increase intratumoral T-cell infiltration in preclinical models ^45^. PLK1 blockade has also enhanced the activity of PD-1/PD-L1 pathway inhibition in preclinical lung and pancreatic tumor models^46,47^. Its recurrent selection by PHAROS is therefore consistent with a second priming mechanism in which attenuation of PLK-driven mitotic programs may render resistant BCC cells more immunologically visible and vulnerable to immune attack released by PD-1 blockade.

To contextualize tumor-intrinsic mechanisms of checkpoint resistance in our cohort, we compiled a gene set from published pan-cancer analyses and reviews of genomic determinants of immunotherapy resistance, focusing on antigen-presentation genes (*B2M*, *HLA-A/B/C*) ^34,48–50^, IFN-γ–JAK–STAT pathway genes (*JAK1*, *JAK2*, *IFNGR1/2*, *STAT1*, *IRF1*, *SOCS1*, and *PTPN2*)^33,34,48,51^, and immune-evasion-associated oncogenic or tumor-suppressor genes (*PTEN*, *MDM2*, *MDM4*, *CTNNB1*, *BRCA2*, *CDK12*)^48–50,52,53^. Mapping these genes onto the JAK1/2 protein–protein interaction network showed substantial overlap between established resistance-associated nodes and targets selected by PHAROS (Fig. 6e). These analyses therefore nominate IFN–JAK–STAT and mitotic/PLK pathways as experimentally testable tumor-cell-priming strategies for PD-1-refractory BCC. The analysis remains exploratory: the BCC states are out of distribution within the Tahoe manifold (Supplementary Fig. S13), and only combination experiments that measure both malignant and immune compartments can determine whether these candidates truly restore sensitivity to PD-1 blockade.

## 3 Discussion

PHAROS addresses an inverse problem that is distinct from standard perturbation-response prediction. Given a source cell-state distribution and a desired target state, it prioritizes drug combinations predicted to move the source toward the target without retraining on the query dataset. Across external combinatorial perturbation datasets, PHAROS recovered exact or mechanism-matched two-drug responses, preserved conversion specificity against random, single-drug, additive, and mismatched-target controls, and extended to cell lines absent from perturbation-model training. Exploratory tumor analyses further showed how patient-derived states can prioritize FDA-approved regimens and nominate pathway-level hypotheses for metastatic breast cancer and immunotherapy-refractory BCC. In BCC, PHAROS identified PLK1- and JAK–STAT-directed candidates that could prime non-responder malignant cells toward a responder-associated state and thereby increase their permissiveness to PD-1-mediated immune killing. Both cohorts were out of distribution; these results are therefore hypotheses for combination testing rather than direct therapeutic predictions. Together, they support PHAROS as a practical framework for screening drug combinations conditioned on specified starting and target cell states.

This inverse-design framework is particularly relevant to oncology, where combination regimens are clinically central but the experimental search space expands rapidly across drugs, doses, schedules, and cellular contexts. Exhaustive single-cell screening therefore remains impractical, even with advances in multiplexed perturbation profiling ^5,18,54,55^. PHAROS is designed to narrow this space by serving as an upfront computational screen that prioritizes combinations for experimental testing according to a specified source-to-target cell-state conversion. The breast-cancer and BCC analyses illustrate complementary use cases: patient-derived states can prioritize existing regimens, nominate previously untested combinations, compare candidates across malignant and immune objectives, and define tumor-cell-priming strategies to be evaluated together with immune-mediated therapies. These examples highlight PHAROS’s key advantage over existing models: it generates data-driven drug-combination hypotheses from specified source and target cell states in primary clinical samples, where matched perturbation data are impractical to obtain. More broadly, the framework could also be used to compare candidate conversion objectives before experimentation by estimating their relative tractability, for example through the predicted reduction in OT distance from baseline. Conceptually analogous to Kolmogorov complexity, this formulation also suggests an operational measure of cell-state complexity: the minimum perturbation sequence required to transform one cellular state into another.

Beyond prioritizing combinations for a single malignant-cell conversion, PHAROS can evaluate the same candidate panel across independently specified objectives in multiple cell populations. In breast cancer, this analysis distinguished regimens with tumor-dominant, immune-supportive, or comparatively balanced predicted profiles, exposing trade-offs that would be obscured by a single malignant-cell endpoint. This multi-objective formulation could be used to prioritize combinations that jointly reverse disease-associated tumor states, restore or preserve favorable immune programs, and avoid undesirable shifts in non-malignant populations. Importantly, these profiles represent parallel cell-type-specific predictions rather than a mechanistic simulation of cell-cell communication or an integrated prediction of clinical efficacy. Their immediate value is therefore in selecting combinations for multicompartment experimental testing.

This creates a practical opportunity for the many primary-tumor and patient-derived single-cell datasets that were collected observationally rather than as perturbation screens. Large cancer atlases now contain tumor and microenvironmental cells across tumor types, clinical states, metastatic sites, and treatment histories^56,57^. PHAROS provides a way to ask therapeutic questions of these data: which approved or repurposable combinations are predicted to move resistant, metastatic, immune-evasive, or stem-like malignant populations toward more desirable reference states while preserving or restoring beneficial immune-cell programs? In this sense, the framework extends single-cell profiling from retrospective biological characterization toward prospective cell-conversion hypothesis generation. Its immediate role is not to replace functional validation, but to reduce a vast combinatorial space to a tractable, state-specific shortlist for testing in organoids, ex vivo cultures, patient-derived models, or prospective clinical studies.

The admissibility framework is central to that use. Rather than treating all query datasets as equally reliable, PHAROS estimates whether the query states are supported by the perturbation-model manifold and whether the requested source-to-target conversion is resolvable. These metrics are not absolute biological truth; they are deployment-risk indicators. The breast-cancer and BCC analyses illustrate the distinction. Out-of-distribution tumor states can still yield clinically coherent hypotheses, but those hypotheses should be labeled as extrapolative and interpreted differently from predictions made within calibrated cell-line support.

PHAROS is also modular. Although implemented here with STATE trained on Tahoe-100M, the search layer can in principle wrap any perturbation model that maps source cells and interventions to predicted post-perturbation states. As single-cell foundation and perturbation models improve, including scGPT, Geneformer, scFoundation, UCE, STATE, and transfer-oriented architectures such as STACK and X-Cell ^7–9,58–61^, PHAROS provides a general route for converting forward models into target-directed virtual screens.

Several limitations define the next stage of development. First, PHAROS inherits the chemical and biological coverage of the perturbation model. In the independent combinatorial perturbation datasets, evaluation was restricted to drug pairs with sufficient overlap with the Tahoe-100M perturbation vocabulary. Second, the current beam search favors trajectories in which intermediate states already move toward the target, which may miss non-monotonic combinations whose benefit emerges only after later perturbations. This can decouple hypothesis-driven modality performance from open-search recovery. A pair may rank highly against random controls in forward scoring yet fail to appear as an exact combination in the retained beam. To tackle this issue, future versions could incorporate global trajectory optimization, reinforcement learning, Monte Carlo tree search, or vector-field objectives such as flux matching to search less greedily through perturbation space ^62^. Third, additional single-cell drug-combination datasets are needed to establish robust benchmarks for models designed specifically to predict combinatorial, rather than single-drug, responses. Fourth, improved transfer to primary tissue will require perturbation training data that better cover tumor lineages, resistance states, clinically approved drugs, and combination regimens. Fifth, the current multi-objective analysis evaluates each cell population independently and therefore does not model cell-cell signaling, compartment-specific drug exposure, systemic toxicity, or emergent interactions within the tumor microenvironment. Future perturbation models trained on multicellular systems, together with combination experiments that jointly measure malignant and immune compartments, will be needed to predict integrated ecosystem responses.

Together, these results show that pretrained perturbation models can be used as therapeutic search engines, not only as response simulators. PHAROS provides one path toward that goal by combining frozen cross-dataset inference, explicit admissibility scoring, and target-directed combinatorial ranking. Its predictions should be treated as prioritized hypotheses, particularly outside the calibrated manifold, but the framework addresses a deployment problem that forward-prediction benchmarks alone do not capture: choosing which combinations to test when only the starting and desired cellular states are known. As perturbation foundation models expand in scale, chemical coverage, and cross-context reliability, this type of modular inverse-design layer could transform single-cell tumor profiles into experimentally actionable, patient-specific combination hypotheses, linking observed disease states directly to the next treatments to test.

## 4 Methods

### 4.1 Single-cell data processing and STATE embedding

All single-cell datasets were processed from raw count matrices. We removed genes detected in fewer than three cells, normalized each cell to a total of 10,000 counts, and applied a log(1 + x) transformation. The processed datasets were embedded with the frozen SE-600M model from STATE (checkpoint se600m_-epoch16.ckpt) using state emb transform (batch size 32); the resulting embeddings were stored in X_state and used for all subsequent quality-control, prediction, and scoring analyses. The embedding model was applied without fine-tuning on any query dataset.

### 4.2 Target calibration of in-domain conversion accuracy

To provide an empirical reference for interpreting target-directed scores, we calibrated single-drug predictions within Tahoe-100M. We selected three cell lines and retained wild-type DMSO controls together with every non-DMSO perturbation measured at 5 µM. For each cell line–drug condition, we applied the matching 5 µM perturbation with the frozen STATE perturbation model to the control population, and compared the predicted and observed treated populations by entropically regularized Sinkhorn optimal transport (OT). We also calculated the control-to-observed-target OT distance and expressed prediction performance as the proportion of this initial distance closed by the predicted transition. These in-domain distributions were used as calibration references for score interpretation, not as universal pass–fail thresholds. Compression and component-selection procedures used for downstream conversion scoring are described separately below.

### 4.3 Embedding-manifold quality control

We assessed whether query cell states were supported by the Tahoe-100M embedding manifold before interpreting a search. The reference comprised all 50 Tahoe cell lines, including DMSO controls and all 5 µM perturbation conditions, with 100 cells sampled per cell line–condition state. In STATE embedding space, we identified the 50 nearest reference cells for each query cell using Euclidean distance and calculated a local-density ratio: the query cell’s mean distance to its neighbours divided by the mean intrinsic neighbour distance of those reference cells. Larger ratios therefore indicate weaker local reference support. We calibrated this statistic using reference cells, whole held-out perturbation states, and held-out cell lines.

At the state level, we called a population OOD when its median local-density ratio was at or above the 99th percentile of the held-out-cell-line calibration. When this calibration was unavailable, we applied the corresponding held-out-state calibration. We also called a population OOD when its held-out-state percentile was at least 99 and at least 25% of its cells exceeded the 99th percentile of the seen-reference-cell distribution, or when at least 50% of its cells exceeded that seen-cell threshold. We called a population boundary when it was not OOD but its held-out-state percentile was at least 95, its seen-reference-state percentile was at least 99, or at least 10% of its cells exceeded the seen-cell 99th percentile. All remaining populations were classified as core. We additionally summarized distance-weighted nearest-neighbour state and tissue annotations to make the biological context of reference support explicit.

### 4.4 Source–target separation quality control

We tested whether each proposed source–target conversion was resolvable in STATE embedding space. Source and target cells were jointly L2-normalized and visualized with UMAP constructed from a cosine-distance neighbourhood graph (15 neighbours; minimum distance 0.3). Separability was quantified independently of the UMAP layout using per-cell k-nearest-neighbour label purity (k = 30, reduced to min(30, n 1) for a population of n cells): for each cell, we calculated the fraction of its non-self neighbours assigned to the same source or target population, then averaged this value within each population. A conversion passed this quality-control gate only when both populations met a mean purity of 0.80; otherwise it was designated poorly resolved and interpreted with additional caution. UMAPs were used for visualization only, whereas the purity criterion determined the separation call.

### 4.5 Conversion-aligned compression for target scoring

For every source–target conversion, we constructed a conversion-aligned linear scoring space with PCA followed by partial least-squares discriminant analysis (PCA–PLS-DA). The projection was fit using the source and target STATE embeddings, after centering and scaling each embedding dimension on the combined fitting data. PCA first reduced the embedding to a candidate number of components, and PLS-DA then identified axes that separated the source and target populations. This transformation was used only to compare predicted and target distributions: the STATE transition model always received and returned embeddings in the original SE space.

We selected the PCA prefilter and PLS-DA dimensions separately for each conversion using PCA candidates of 96, 128, 192 and 256 and PLS-DA candidates of 64, 96, 128 and 192 components. For conversions with at least 150 cells in each population, we performed 10 repeated stratified splits, fitting each candidate projection on 50% of the source and target cells and evaluating normalized source–target centroid separation in the held-out cells. We selected the least complex candidate, that is the one with the lowest values for PCA and PLS-DA parameters, whose mean held-out separation was within one standard error of the best candidate, that is the one maximizing centroid separation in the held-out cells. For smaller populations, for which this selection procedure was not stable, we used the prespecified fallback of 128 PCA and 64 PLS-DA components. The final projection was subsequently fit using the automatic fit/evaluation policy: source and target cells were held out for scoring when both populations contained more than 512 cells and otherwise were fit and scored using all available cells. Projected components were not whitened, thereby retaining their fitted relative variance.

All predicted and observed populations were projected with the conversion-specific map before target scoring. Because linear projection changes the scale and geometry of the embedding, we did not L2-normalize projected embeddings, used squared Euclidean costs for Sinkhorn OT, and set the entropic regularization parameter to 0.1 times the median pairwise cost within the projected target population. Thus, compression focused the comparison on variation that distinguished the requested conversion without modifying the state transitions being evaluated.

### 4.6 Shared PHAROS conversion engine

Both hypothesis-driven evaluation and open search used the same frozen inference engine. For each requested conversion, source and target cells were identified from a state-label column in the embedded AnnData object and their SE embeddings were loaded from X_state. We evaluated five independently sampled source–target batches per conversion, except in the high-sensitivity analysis described below, which used three selected batches. Each source and target batch contained 256 cells when available; when a population contained fewer than 256 cells, we used batches of 128 cells, and when it contained fewer than 128 cells, we used batches of 64 cells. Sampling with replacement was used only when explicitly permitted and the available population was smaller than the requested batch. The unperturbed source population provided the common baseline for each conversion.

We loaded the Tahoe ST-SE state-transition model from the pretrained checkpoint located in the state_generalization_X_state directory and used it without retraining or fine-tuning. The model accepts a population of SE embeddings and a perturbation label from its trained Tahoe vocabulary and returns one predicted SE embedding per input cell. For an ordered path of perturbations (p_1_, …, p_L_), we recursively applied the transition model, S_t+1_ = F_θ_(S_t_, p_t+1_), beginning with the observed source population S_0_. Each successive drug was therefore evaluated in the state predicted after the preceding drug, preserving cell-level correspondence through the path. This state-dependent composition differs from an additive-vector approximation, in which single-drug displacements are estimated from the same initial state and summed, and allows the predicted result to depend on treatment order.

Formally, for a path p = (p_1_, …, p_L_), the recursively predicted terminal population is S^^^p = F_θ_(… F_θ_(S_0_, p_1_), …, p_L_). Let Π denote the conversion-specific scoring map, which is the identity in uncompressed analyses and the fitted PCA–PLS-DA map when compression is used. We defined the target-distance objective as

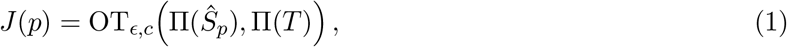

where T is the observed target population and OT_ɛ,c_ is the entropically regularized Sinkhorn OT distance under the specified cell–cell cost c. All populations were represented as uniformly weighted empirical distributions. Lower J(p) indicates a closer predicted match to the target. Relative conversion performance was summarized as the fraction of the unperturbed source–target distance closed,

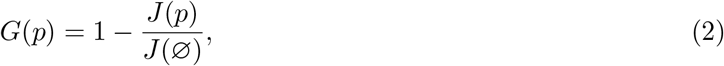

where J(∅) = OT_ɛ,c_(Π(S_0_), Π(T)) is the baseline distance without an applied perturbation. Candidate generation differed between the two analysis modes, but all candidates were propagated and scored with these same frozen transition and distribution-scoring procedures.

### 4.7 High-sensitivity batch selection

We applied high-sensitivity batch selection only to source–target conversions that failed the source– target separation quality-control criterion. Its purpose was to identify locally coherent source and target subpopulations with stronger baseline separation, thereby enabling a sensitivity analysis when the complete populations could not be resolved reliably. For each such conversion, we generated 1,000 candidate batch pairs by selecting a random seed cell independently from the source and target populations and forming each batch from its nearest neighbours in STATE embedding space. Batch sizes followed the 256-, 128-, or 64-cell schedule described above.

We scored every candidate pair by its unperturbed source-to-target Sinkhorn OT distance in the conversion-specific scoring space and retained the three pairs with the largest baseline distance. We used no overlap penalty (–batch-overlap-penalty 0); thus, selection prioritized maximal source– target separation without penalizing cell reuse across the retained batches. The selected batches replaced random sampling for both hypothesis-driven and open-search analyses of these failed-separation conversions. Because this procedure conditions the evaluation on locally separable subpopulations, high-sensitivity results quantify performance in informative subsets rather than across the full source and target populations.

### 4.8 Hypothesis-driven combination evaluation

In the hypothesis-driven mode, we evaluated a user-defined set of candidate interventions against a specified source–target conversion. Each conversion included 100 random two-drug control pairs and used a Sinkhorn OT metric specified as cosine distance with entropic regularization ɛ = 0.05 and 100 Sinkhorn iterations. When the conversion-aligned projection was active, scoring followed the projected-space metric and regularization procedure described above. Candidate order and dose were selected on the first source–target batch; the selected candidates were then evaluated across all five batches, or across the three high-sensitivity batches where applicable.

For every candidate two-drug combination, we enumerated both treatment orders and all concentrations represented for each constituent drug in the Tahoe perturbation-model vocabulary. We sequentially predicted every order–concentration configuration, calculated its Sinkhorn OT distance to the target, and retained the configuration with the lowest distance. This procedure was applied equivalently to explicit candidate pairs, mechanism-of-action (MOA)-matched pairs, and random controls, so that comparisons were not driven by an arbitrary treatment order or dose choice. For pairs defined by MOA category, drug identities were matched against the fine-grained MOA annotations; MOA terms were oriented consistently with the selected explicit-pair order when an explicit pair was supplied.

The workflow supported several hypothesis-driven use cases. First, a fully specified two-drug pair tested whether a nominated combination was predicted to reach the target state. Second, a single named drug paired with one MOA term tested that drug against all vocabulary-compatible partners sharing the specified mechanism, enabling mechanism-based substitution when one constituent drug was absent from the model vocabulary. Third, two MOA terms defined a broader set of candidate pairs spanning the two requested mechanism classes. Finally, panel analysis accepted multiple named pairs and/or categories, selected the optimal order and concentration for each panel member, and evaluated all members with the same source–target populations and scoring procedure, enabling direct comparison of clinically or mechanistically distinct treatment hypotheses.

We compared the distributions of per-batch candidate-pair Sinkhorn OT values with those of the 100 random pairs using one-sided Mann–Whitney U tests, with the alternative hypothesis that random-pair OT values were greater than those of the nominated group. For each conversion, tests compared random pairs with MOA-matched pairs and, when supplied, with the explicit pair. We adjusted P values across all comparisons displayed in each multi-conversion report using the Benjamini–Hochberg procedure and reported the resulting false-discovery-rate-adjusted Q values.

### 4.9 Open combinatorial search

For open search, PHAROS used diverse beam search to explore ordered two-drug paths without specifying candidate pairs in advance. Starting from the observed source population, we considered all non-control perturbation labels in the Tahoe vocabulary, excluding reuse of a base drug within the same path even when a different concentration label was available. We used a maximum depth of two, retained a beam of 128 paths at each depth, and processed perturbation labels in chunks of 16. Thus, each depth extended every retained partial path by all allowable next perturbations and evaluated the resulting predicted populations against the requested target.

We used Sinkhorn OT throughout the search rather than an energy-distance prefilter. At each depth, candidate paths were first ranked with a 10-iteration Sinkhorn prefilter, after which the best 10 × 128 = 1,280 candidates were reranked using the full Sinkhorn calculation (cosine metric specified at the command level, ɛ = 0.05, 100 iterations; projected-space settings are described above). The 128 paths retained after reranking were selected greedily with a path-overlap penalty of 25. Specifically, for each candidate, we added 25 times its maximum fractional overlap in base-drug identities with any already retained path to its ranking score. This encouraged the beam to preserve mechanistically distinct alternatives instead of filling with paths that differed only trivially from the leading prediction.

To prioritize paths that generalized across sampled cells, we robustly reranked the full-Sinkhorn candidate pool at every search depth. Each candidate path was replayed from five independently sampled source populations and compared with the corresponding target populations using Sinkhorn OT. For B = 5 batches, we ranked candidates by

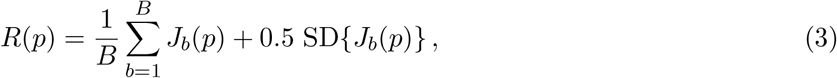

where J_b_(p) is the target-distance objective for batch b. Thus, lower R(p) favoured both low predicted target distance and reproducibility across batches. The diverse-beam overlap penalty was applied after this robust reranking. The final output therefore comprised ordered paths selected for target proximity, cross-batch robustness, and diversity relative to other retained paths.

### 4.10 Target-pair recovery against a beam-matched Monte Carlo null

We quantified the statistical significance of open-search recovery using a per-conversion Monte Carlo null model that matched the size and basic constraints of the depth-two beam. For each conversion, we examined the 128 depth-two paths retained by the search, ranked by Sinkhorn OT. Drug names were normalized before matching to the experimentally defined target pair, including removal of salt and formulation terms. We recorded the total number of appearances of each target constituent across all retained path positions, whether the unordered target pair occurred together in at least one path, and, when it did, the best Sinkhorn rank of an exact-pair path.

The null model represented a beam with no target-directed signal while preserving the Tahoe search-space structure. In each Monte Carlo replicate, we sampled 128 distinct first-step perturbation labels without replacement from a universe of 379 drugs at three concentrations per drug. We then sampled 128 unique ordered two-step label paths by selecting a first-step label from this set and a second-step label uniformly from the legal extensions. By default, legal extensions excluded all concentrations of the same base drug, matching the search constraint that a drug cannot be reused within a path. The simulated beam therefore preserved the beam width, concentration multiplicity, ordered-path structure, and no-repeated-drug rule, but assigned drugs independently of the target and predicted scores.

For each target constituent and for exact-pair recovery, we generated 10,000 simulated beams and calculated an upper-tail empirical P value as 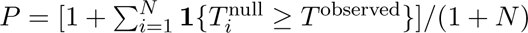, where **1**{·} denotes the indicator function, T is the relevant recovery statistic, and N is the number of simulations. Exact-pair P values were reported only when the pair was observed in the retained beam. This analysis tests whether enrichment of either target drug, or recovery of the complete target pair, exceeded that expected from a random beam under the same nominal search-space constraints.

### 4.11 Metastatic-to-primary reference matching in the breast-cancer analysis

The primary and metastatic malignant-cell samples in the breast-cancer dataset were obtained from different individuals; consequently, no metastatic sample had a patient-matched primary tumor reference. To define a consistent conversion objective across metastatic samples without selecting a primary reference arbitrarily, we matched each metastatic sample to the transcriptionally closest primary malignant-cell sample before drug-conversion analysis. This matching was used solely to standardize the source–target task; it was not interpreted as evidence of tumor lineage, evolutionary ancestry, or a patient-specific clinical relationship.

We performed matching using the frozen STATE embeddings. For each metastatic sample and each candidate primary sample, we drew 256 malignant cells per sample for each of 100 bootstrap replicates, sampling with replacement when a sample contained fewer than 256 cells. We L2-normalized embeddings and calculated the uniformly weighted, cosine-cost Sinkhorn OT distance between every metastatic– primary batch pair (ɛ = 0.05; 100 iterations). For each metastatic–primary pair, we summarized bootstrap distances by their mean plus 0.5 times their standard deviation; the primary sample with the lowest summary score was selected as the reference target. We also recorded the frequency with which each primary reference won across bootstrap replicates, reported the three best-scoring primary candidates, and performed pooled-primary scoring as a sensitivity analysis. Target matching was completed before conversion screening and did not use intervention identity, drug approval status, or predicted conversion performance.

### 4.12 Cross-cell-type objective profiling in metastatic breast cancer

To compare treatment profiles across malignant and lymphoid populations, we restricted the breast-cancer atlas to HR^+^/HER2^−^ tumors and pooled cells across patients because individual samples contained insufficient numbers of each lymphoid state for patient-level analysis. Cell identities were taken directly from the published minor_celltype annotations. We defined six source-to-target objectives: metastatic-to-primary malignant cells; metastatic-to-primary GZMK^+^ CD8^+^ effector-memory cells; metastatic-to-primary ZNF683^+^ tissue-resident-memory CD8^+^ cells; metastatic CXCL13^+^ exhausted CD8^+^ cells to primary ZNF683^+^ tissue-resident-memory CD8^+^ cells; metastatic-to-primary CD16^+^ NK cells; and metastatic-to-primary memory B cells.

Each objective was evaluated in the hypothesis-driven mode using the same panel of nine FDA-approved two-drug combinations and 100 random two-drug controls, with the shared inference, projection, order-and-concentration selection, and Sinkhorn OT settings described above. Malignant and exhausted-CD8-to-resident-memory objectives used five randomly sampled source–target batches. The GZMK^+^ effector-memory, ZNF683^+^ resident-memory, CD16^+^ NK-cell, and memory-B-cell objectives used the high-sensitivity procedure described above, with three batches selected from 1,000 candidate batch pairs. Batches contained 256 cells for the malignant and GZMK^+^ objectives and 128 cells for the remaining objectives. For each conversion, the first batch was used to select treatment order and concentration and was excluded from the reported performance summaries. Random-control scores were first averaged across the remaining evaluation batches at the drug-pair level. We then calculated the empirical superiority percentile of each FDA-approved combination as the percentage of random pairs with an equal or greater mean Sinkhorn OT distance to the target; higher values therefore indicated more favorable target-state alignment. The six objectives were modeled independently and were used to compare cross-cell-type treatment profiles rather than to simulate interactions among cellular compartments.

### 4.13 Analysis of malignant cell conversion for basal cell carcinoma

We downloaded the single-cell BCC dataset from the 3CA Cancer Cell Atlas (https://www.weizmann.ac.il/sites/3CA/skin), which annotates malignant cells by copy-number-alteration inference^56^. We retained malignant cells from post-immunotherapy biopsies only. Clinical anti-PD-1 response was obtained from Supplementary Table 1 of Yost et al. 2019^31^. Because of the limited number of malignant cells per patient, we pooled cells from responders and non-responders separately. The pooled post-treatment non-responder population was defined as the source and the pooled post-treatment responder population as the target. This analysis therefore tests a tumor-cell-priming objective: identification of perturbations predicted to shift malignant cells from a non-responder-associated state toward a responder-associated state, to be evaluated in combination with PD-1 blockade rather than as a model of the immune response itself.

The pooled populations passed the source–target separation admissibility check. We therefore ran PHAROS in open-search mode using the default settings described above. The out-of-distribution assessment was retained as a separate admissibility result and was used to designate the analysis as exploratory.

For the network analysis, we queried the Homo sapiens JAK2 protein in the full STRING network on August 8, 2026, retaining interactions with the highest confidence threshold (0.9) and limiting the network to 20 interactions. Edges were weighted by STRING’s experimental-evidence score and displayed using a deterministic Fruchterman–Reingold layout. For annotation of the network, nodes encoded by JAK or STAT family members were designated as targets of the drugs categorized as JAK–STAT inhibitors.

## 5 Data availability

All datasets used in this work are publicly available. Tahoe-100M can be downloaded from https: //huggingface.co/datasets/tahoebio/Tahoe-100M. The A549 dataset is accessible from GEO under GSE206741. The sciPlex dataset of three brain cancer cell lines is accessible from GEO under GSM7056151. The metastatic breast cancer dataset is accessible from https://zenodo.org/records/13743374. The pre-processed basal cell carcinoma dataset can be downloaded from the 3CA cancer single-cell atlas https: //www.weizmann.ac.il/sites/3CA/skin. Processed datasets used to generate results are available at https://zenodo.org/records/21925263.

## 6 Code availability

PHAROS software and tutorials are available under https://github.com/jbezney61/PHAROS. All code and scripts used to reproduce the work in this paper are available under https://github.com/jbezney61/PHAROS_reproduce.

## Acknowledgements

We thank Andrea Celli and members of the CompBio Lab at Bocconi University for helpful discussions and feedback throughout the development of this work. We also thank members of the Steinmetz lab for discussions around the theoretical work and motivation behind this paper. We thank Jesus Miguens Blanco and Rogelio Hernandez-Lopez for discussions around applications in oncology and inspiration for the biological importance of combinatorial perturbations.

Common AI tools were used to improve the clarity and readability of the manuscript. The authors reviewed and edited all generated content and take full responsibility for the final publication.

## Ethics declaration

L.M.S is co-founder and shareholder of Sophia Genetics, Recombia Biosciences, and LevitasBio. L.S.Q. is a founder of Epicrispr Biotechnologies. All other authors declare they have no competing interests.

## Supplementary Information

### S1 Supplementary Figures

**Figure S1:**
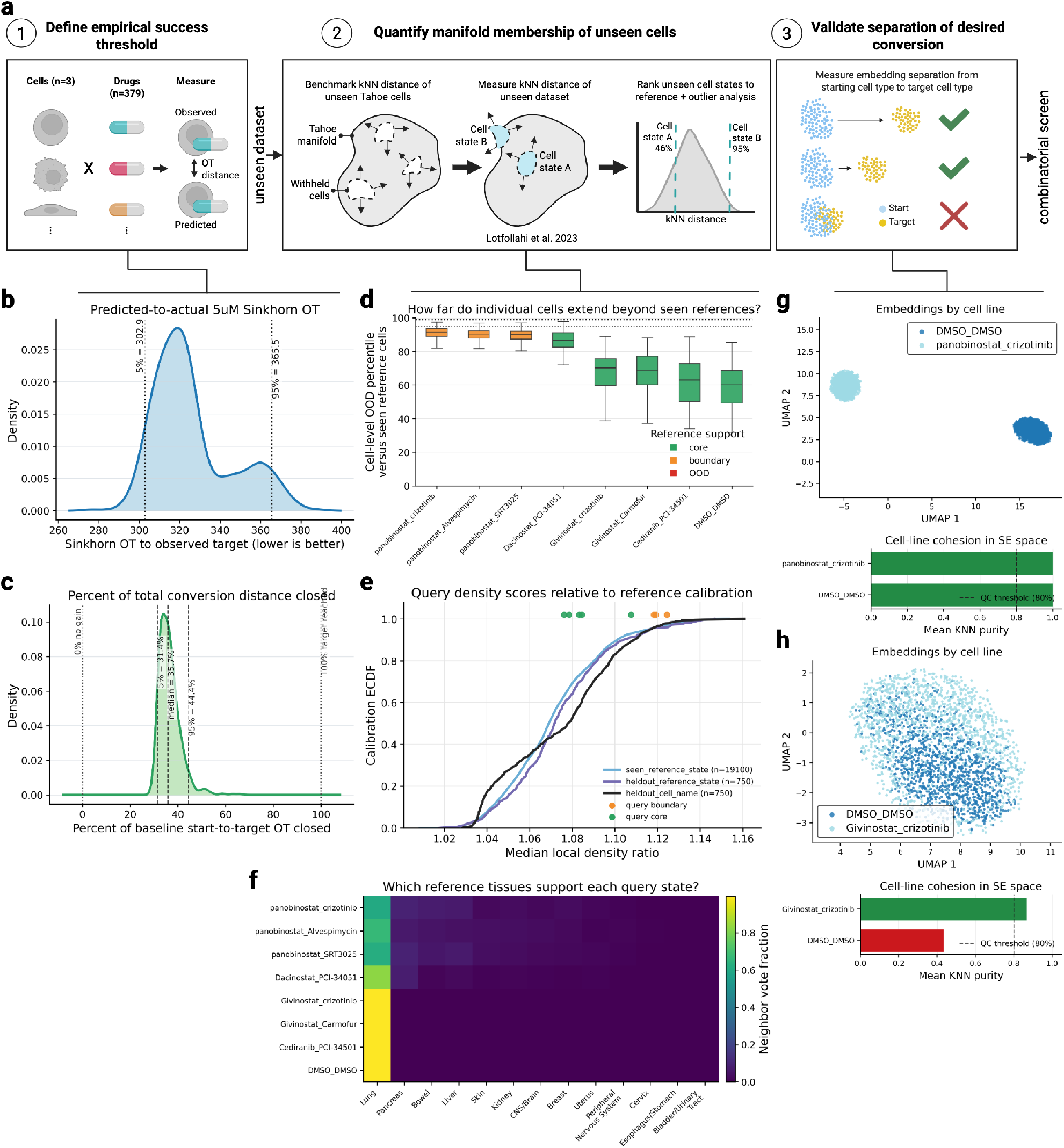
Admissibility framework. **a,** Schematic of 3-part admissibility framework for unseen data. **b,** Distribution of sinkhorn OT distance from predicted single drug perturbation to observed perturbations across 3 cell lines and all 379 drugs at 5uM within Tahoe-100M. **c,** Distribution of source-to-target OT distance closed across 3 cell lines and all 379 drugs at 5uM within Tahoe-100M. **d,** Cell-level manifold percentiles relative to withheld Tahoe-100M cells across eight, 2-drug perturbations from unseen A549 data. Query states categorized into core (green), boundary (yellow), and out of distribution (red). **e,** Empirical cumulative distributions show median local-density ratios for seen, withheld-state, and withheld-cell-line Tahoe-100M calibrations. Larger values indicate weaker reference support. Query states are plotted above the distributions and colored by their support classification. **f,** Distance-weighted tissue annotations of the nearest Tahoe-100M reference neighbors for each query state. Rows denote query states, columns denote reference tissues and color indicates the fraction of total neighbor weight assigned to each tissue. **g,** An example of a specific conversion from start (DMSO) to target (panobinostat + crizotinib) that passed separation. Top: UMAP in SE embedding space. Bottom: barplot of mean KNN purity per cell state. **h,** An example of a specific conversion from start (DMSO) to target (givinostat + crizotinib) that failed separation. Top: UMAP in SE embedding space. Bottom: barplot of mean KNN purity per cell state.

**Figure S2:**
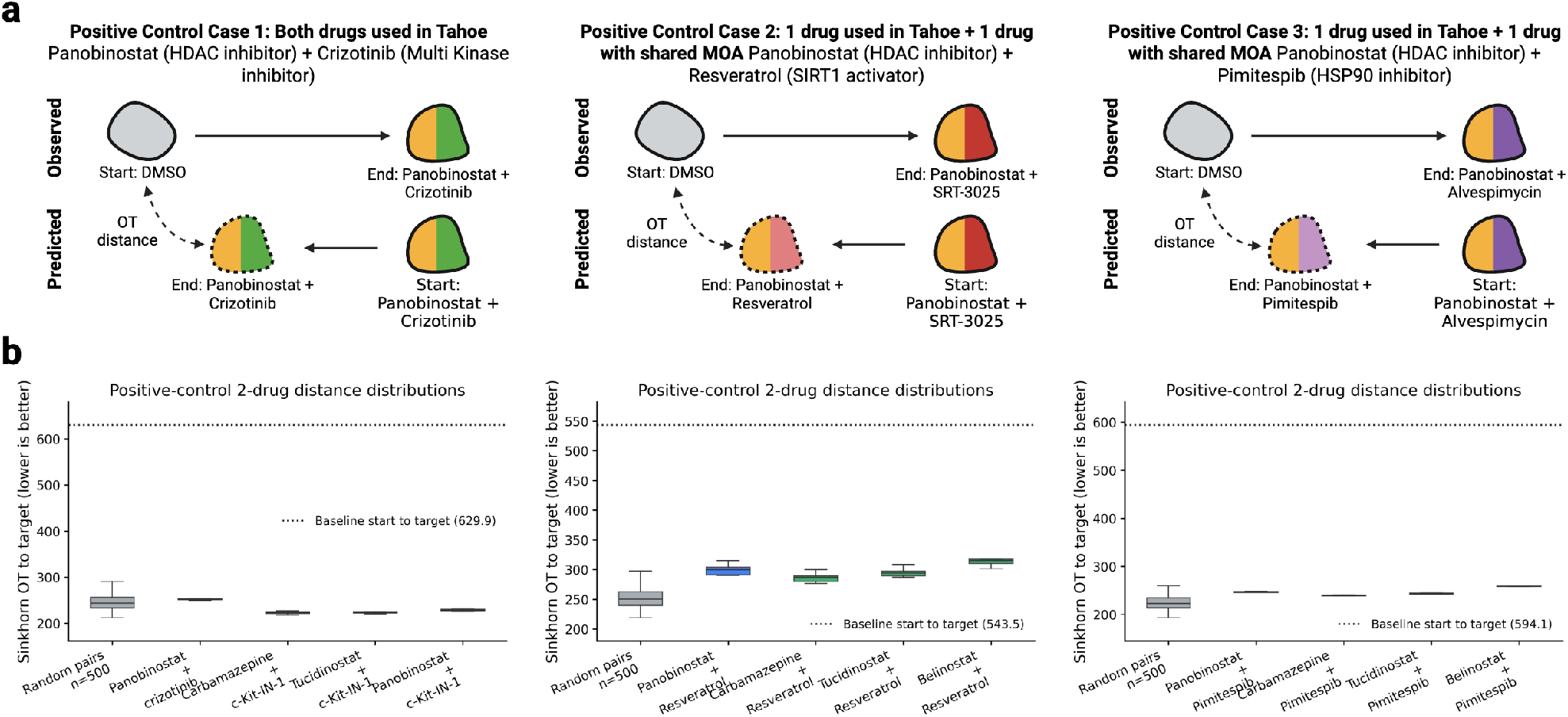
Efficacy when start and target are reversed. **a,** Schematic showing 3 positive control cases where the starting cell distribution and target cell distribution are reversed during prediction. In PHAROS the starting cell state is the DMSO no-drug state, the prediction is the change in latent space given the 2-drugs, and the prediction is compared against the experimentally observed 2-drug state. In this reversed orientation, the starting cell state is the experimentally observed 2-drug state, the prediction is the change in latent space given the 2-drugs, and the prediction is compared against the DMSO no-drug state. **b,** Distribution of sinkhorn OT distance to target under 2-drug predicted perturbations comparing 100 random pairs (grey), the exact 2-drug pair (blue), and all pairs with the same MOA (green). For each unordered drug pair, we evaluated all tested concentration combinations and both treatment orders, and retained the configuration with the lowest OT distance to the target. All 2-drugs were ran across 5 batches of starting and target cells. Dotted line is baseline OT distance between starting cell state and target cell state.

**Figure S3:**
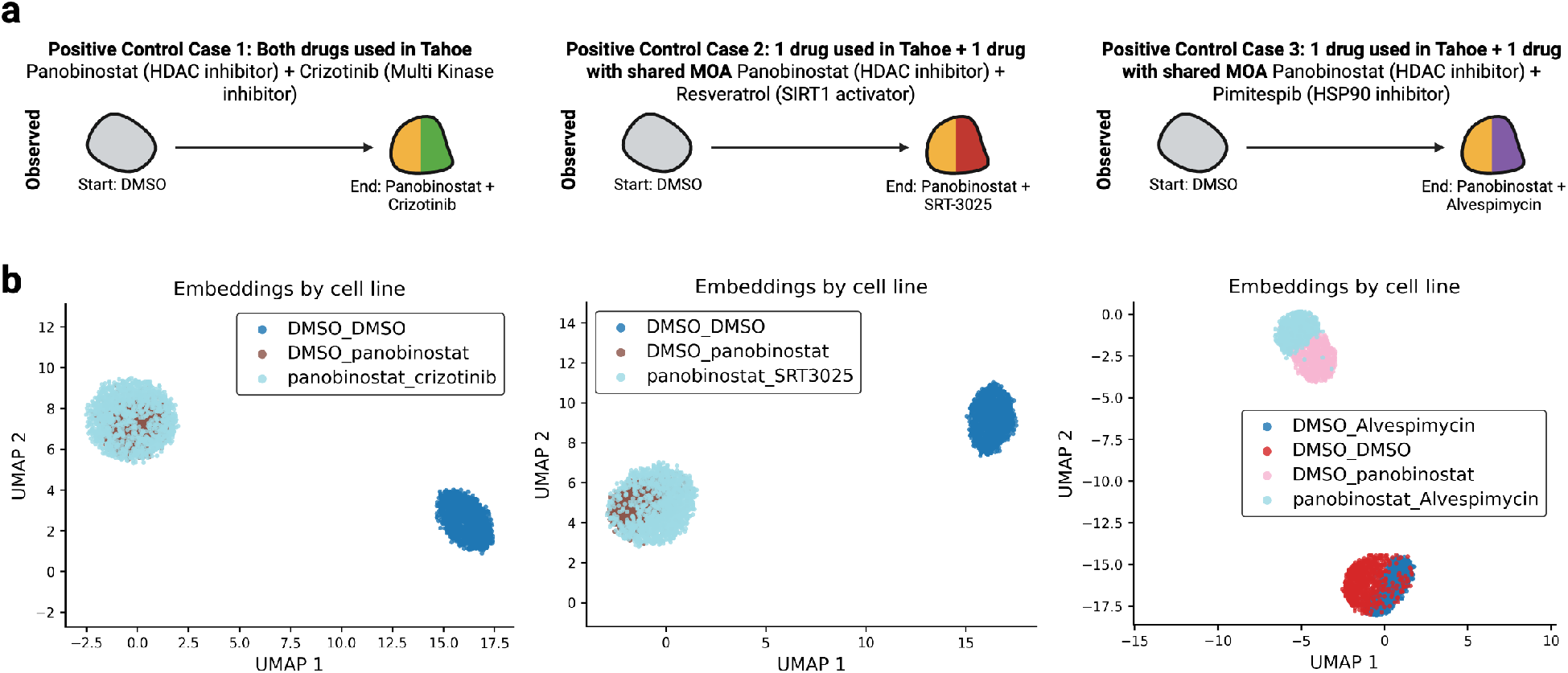
UMAP separation of experimental conditions in STATE single-cell foundation model embedding space. **a,** Schematic showing 3 positive control cases of 2-drug perturbations. **b,** UMAP visualization of experimentally measured cell states relevant to each of the 3 positive control cases. Case 1 and 2 contained just one of the single drug perturbations (DMSO + panobinostat), while case 3 contained both single drug perturbations.

**Figure S4:**
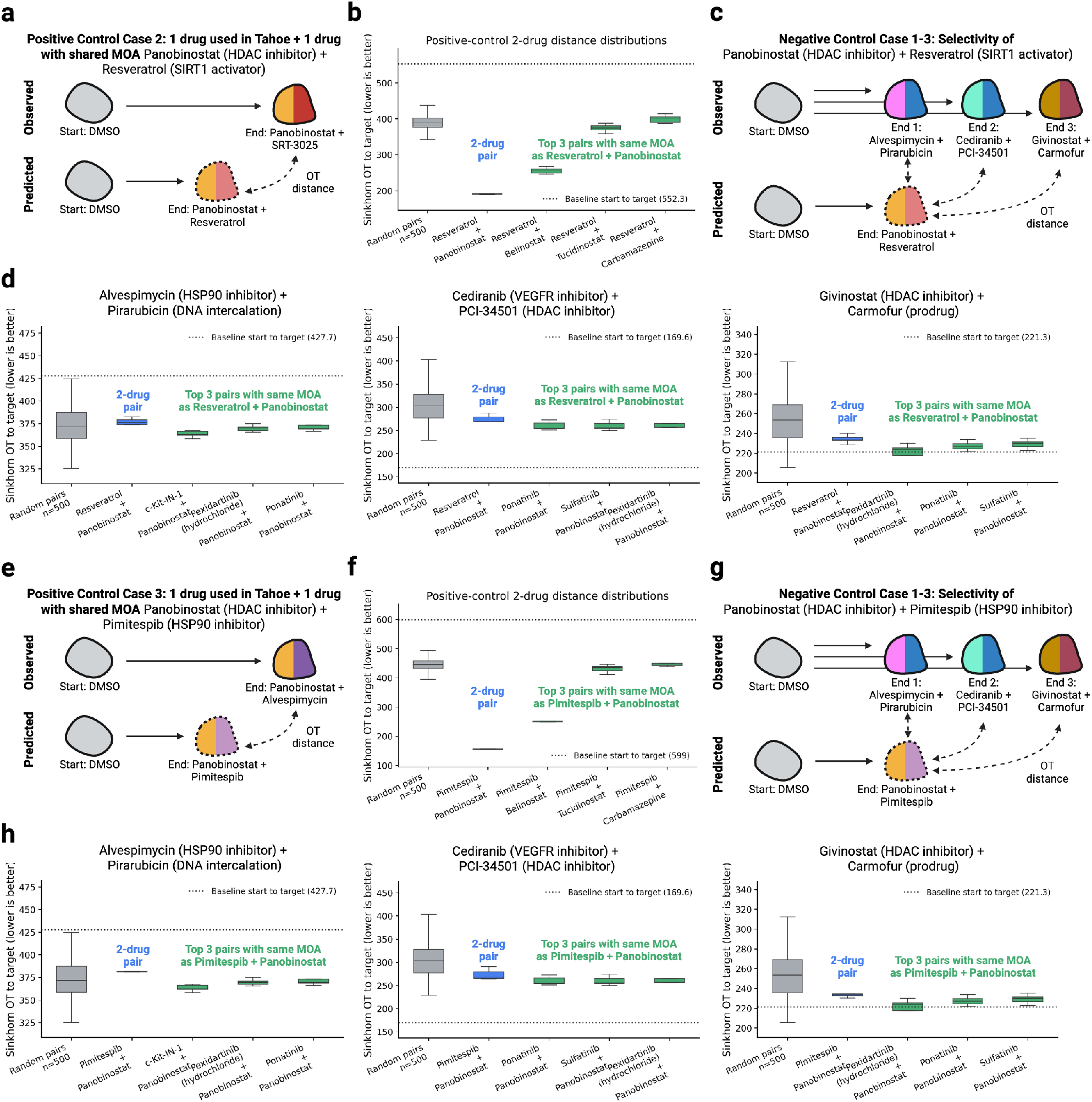
Specificity of positive control cases 2-3. **a,** Schematic of observed versus predicted for positive control case 2. **b,** Observed matches predicted. Distributions of OT distance to target comparing 100 random 2-drugs (grey), the panobinostat + resveratrol mechanism-matched proxy pair (blue), and top 3 pairs with matching MOA (green). **c,** Schematic demonstrating 3 counter-factual examples where the predicted pair, panobinostat + resveratrol, is compared against the observed pairs of alvespimycin + pirarubicin, cediranib + PCI34501, and givinostat + carmofur. **d,** 3 scenarios where observed does not match predicted. Distributions of OT distance to target comparing 100 random 2-drugs (grey), the panobinostat + resveratrol mechanism-matched proxy pair (blue), and top 3 pairs with matching MOA (green). **e,** Schematic of observed versus predicted for positive control case 3. **f,** Observed matches predicted. Distributions of OT distance to target comparing 100 random 2-drugs (grey), the panobinostat + pimitespib mechanism-matched proxy pair (blue), and top 3 pairs with matching MOA (green). **g,** Schematic demonstrating 3 counter-factual examples where the predicted pair, panobinostat + pimitespib, is compared against the observed pairs of alvespimycin + pirarubicin, cediranib + PCI34501, and givinostat + carmofur. **h,** 3 scenarios where observed does not match predicted. Distributions of OT distance to target comparing 100 random 2-drugs (grey), the panobinostat + pimitespib mechanism-matched proxy pair (blue), and top 3 pairs with matching MOA (green).

**Figure S5:**
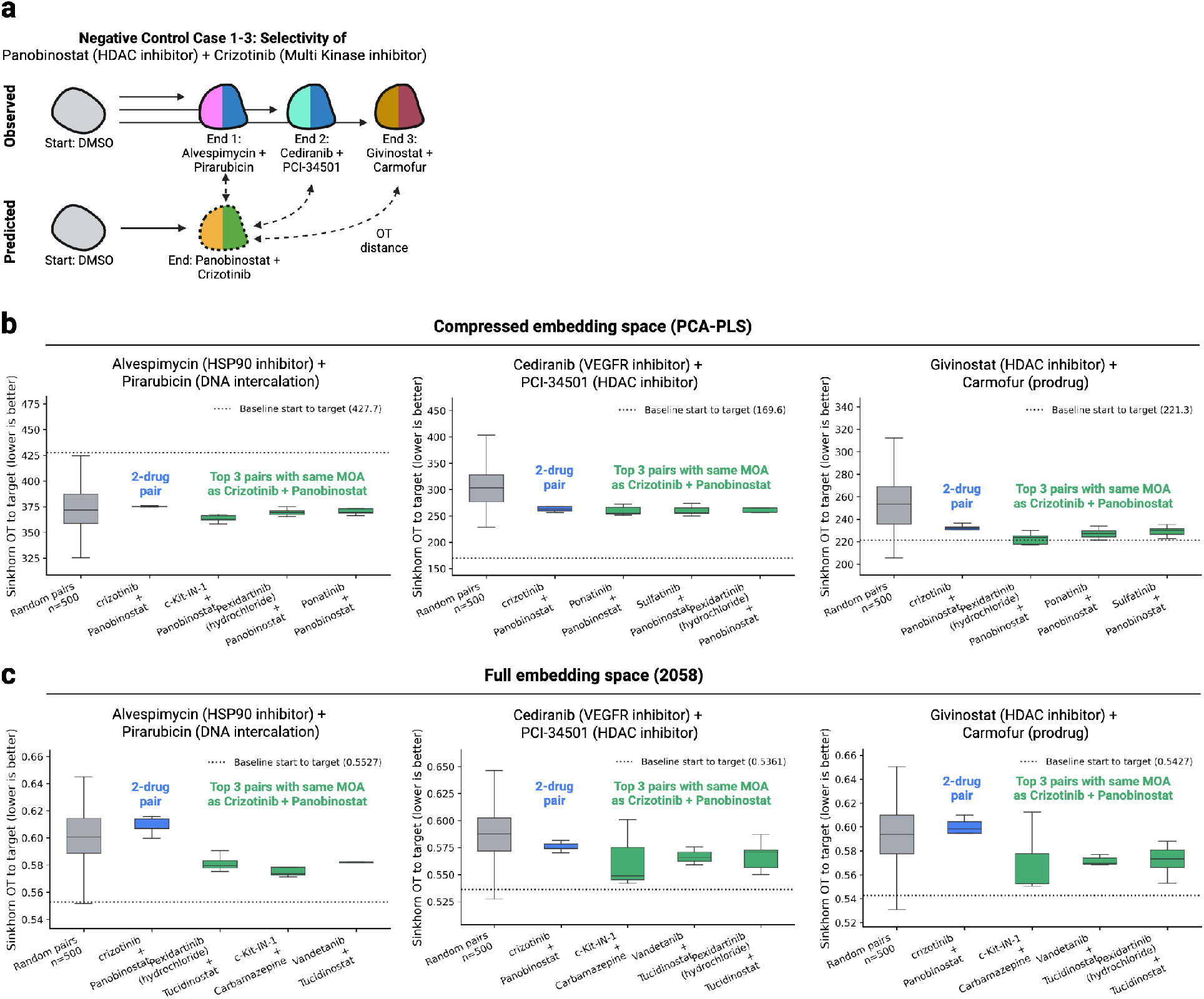
Effect of compression on specificity. **a,** Schematic demonstrating 3 counter-factual examples where the predicted pair, panobinostat + crizotinib, is compared against the observed pairs of alvespimycin + pirarubicin, cediranib + PCI34501, and givinostat + carmofur. **b,** Results shown in Fig 3d. Generated with default PCA-PLS compression. 3 scenarios where observed does not match predicted. Distributions of OT distance to target comparing 100 random 2-drugs (grey), exact panobinostat + crizotinib (blue), and top 3 pairs with matching MOA (green). All 2-drug predictions were evaluated across 5 batches of starting and target cells. **c,** 3 scenarios where observed does not match predicted. Generated without PCA-PLS compression, operating in full dimensionality (2058). Distributions of OT distance to target comparing 100 random 2-drugs (grey), exact panobinostat + crizotinib (blue), and top 3 pairs with matching MOA (green). All 2-drug predictions were evaluated across 5 batches of starting and target cells.

**Figure S6:**
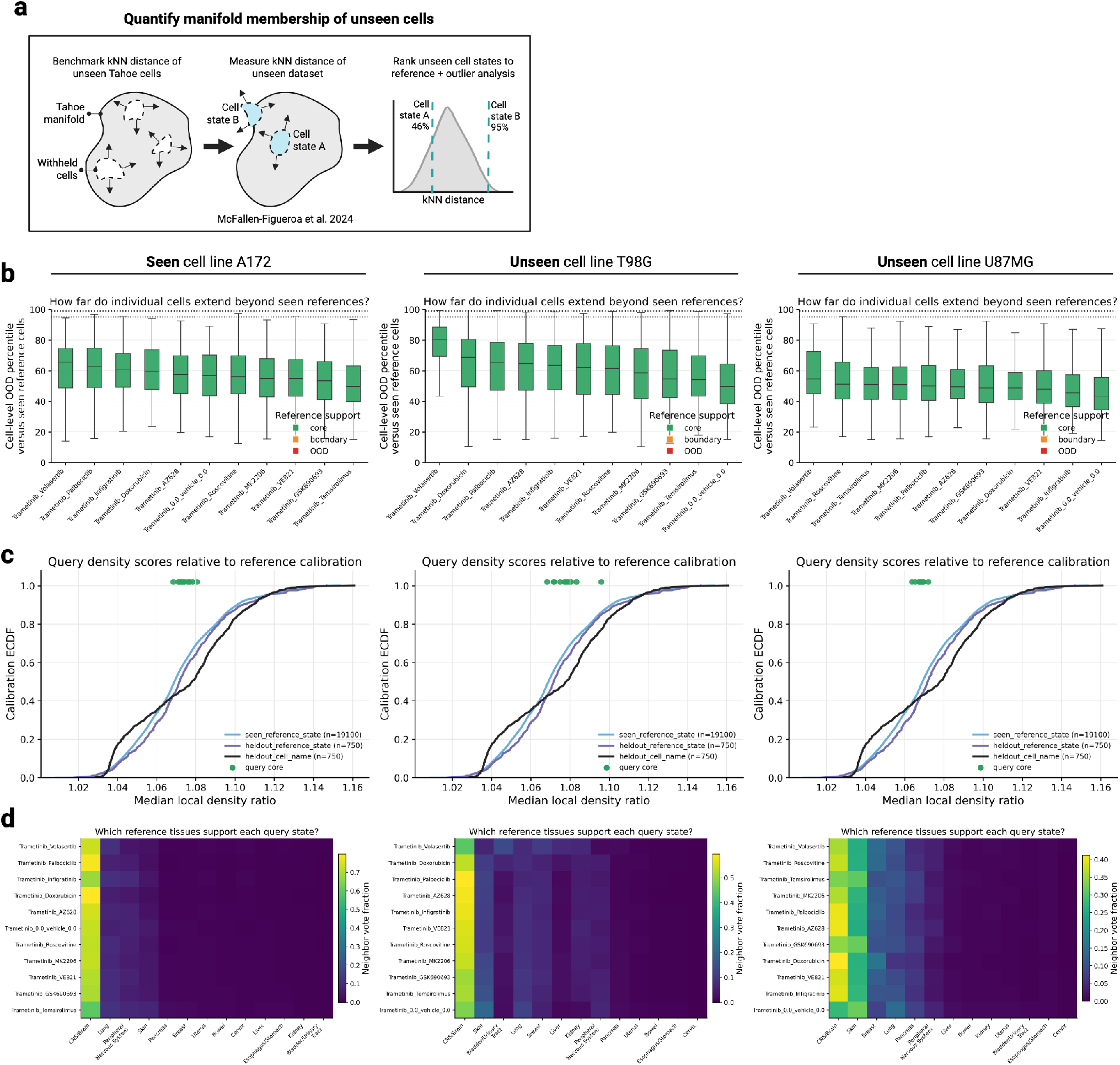
Manifold admissibility of 3 brain cancer cell lines. **a,** Part 2 of the admissibility framework. Quantify Tahoe-100M manifold membership of unseen cell states. **b-d,** Left: seen cell line A172. Middle: unseen cell line T98G. Right: unseen cell line U87MG. **b,** Cell-level manifold percentiles relative to withheld Tahoe-100M cells. Query states categorized into core (green), boundary (yellow), and out of distribution (red). **c,** Empirical cumulative distributions show median local-density ratios for seen, withheld-state, and withheld-cell-line Tahoe-100M calibrations. Larger values indicate weaker reference support. Query states are plotted above the distributions and colored by their support classification. **d,** Distance-weighted tissue annotations of the nearest Tahoe-100M reference neighbors for each query state. Rows denote query states, columns denote reference tissues and color indicates the fraction of total neighbor weight assigned to each tissue.

**Figure S7:**
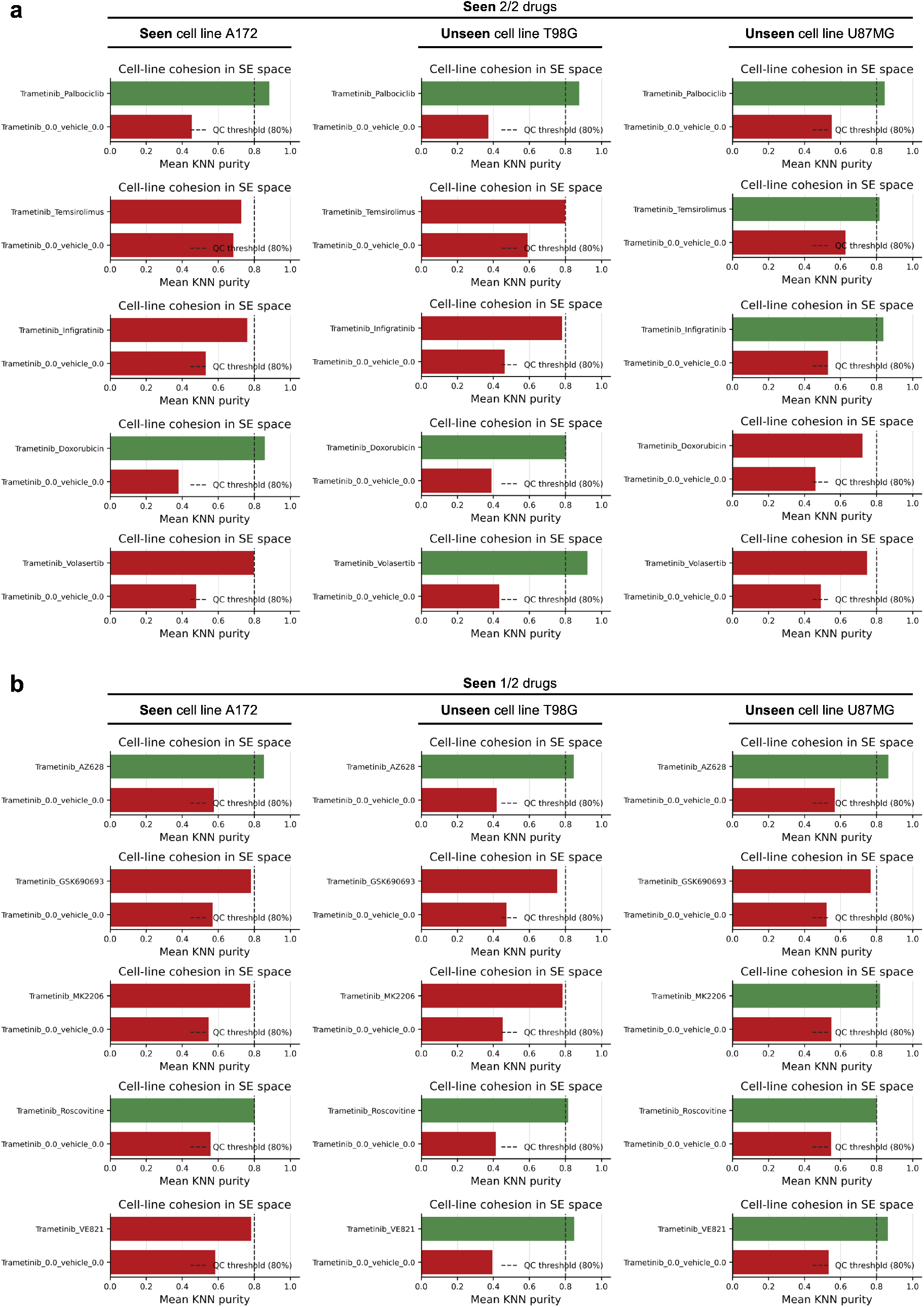
Source-target separation admissibility for every conversion within 3 brain cancer cell lines. **a-b,** Left: seen cell line A172. Middle: unseen cell line T98G. Right: unseen cell line U87MG. **a,** Part 3 of the admissibility framework for seen 2/2 drugs. Barplots of mean KNN purity per cell state for every conversion. **b,** Part 3 of the admissibility framework for seen 1/2 drugs. Barplots of mean KNN purity per cell state for every conversion.

**Figure S8:**
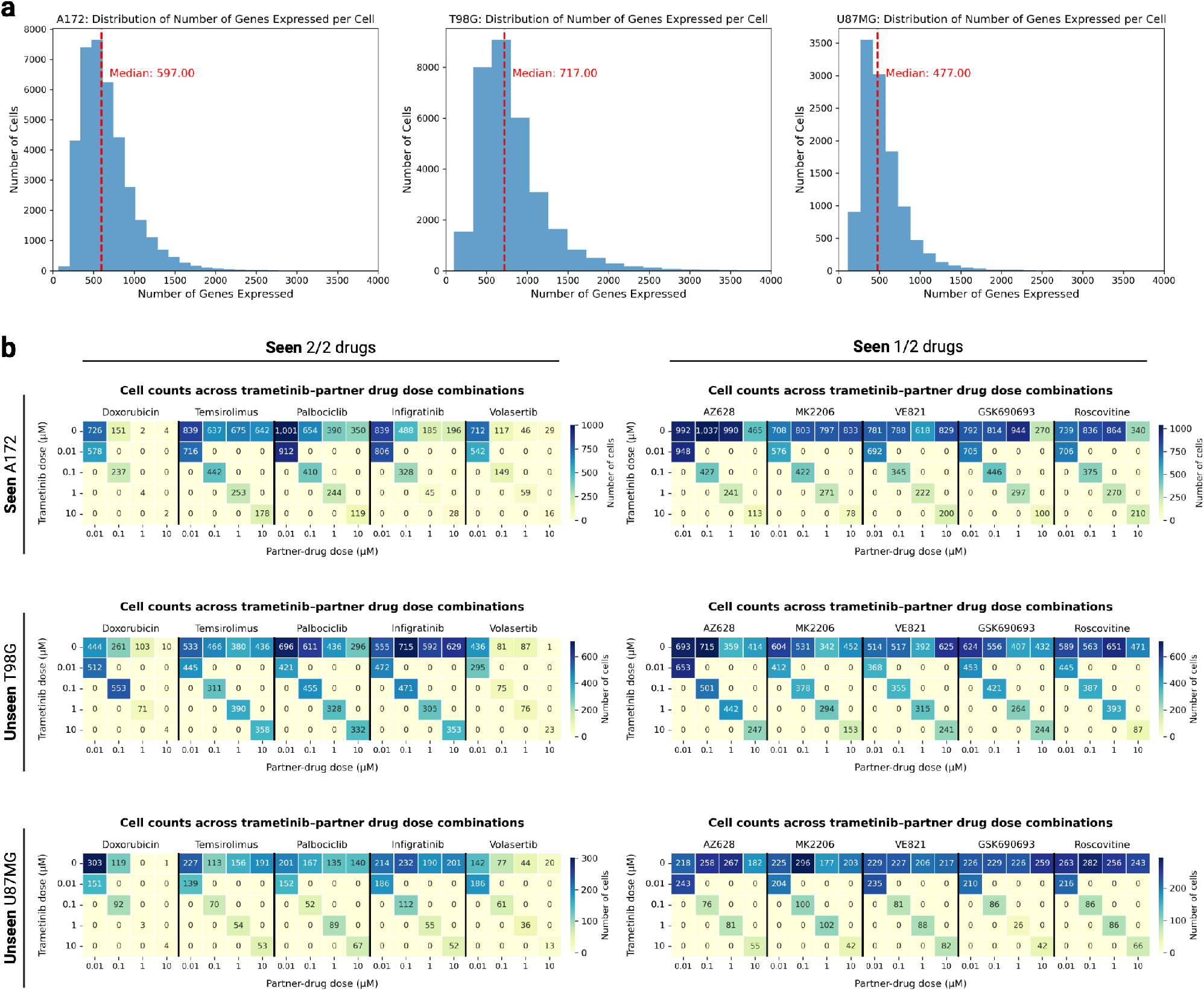
QC metrics for the 3 brain cancer cell lines. **a,** Distribution of genes captured per cell for seen A172 (left), unseen T98G (middle), and unseen U87MG (right). Red dotted line indicates median value. **b,** Left: seen 2/2 drugs. Right: seen 1/2 drugs. Top: seen cell line A172. Middle: unseen cell line T98G. Bottom: unseen cell line U87MG. Heatmap of total number of cells found in each 2-drug perturbation condition. Y-axis denotes the five concentrations (uM) of trametinib (0, 0.01, 0.1, 1, 10), which is paired with every other drug. X-axis denotes the five other paired drugs separated by a black line and categorized into their respective 4 concentrations in uM (0.01, 0.1, 1, 10).

**Figure S9:**
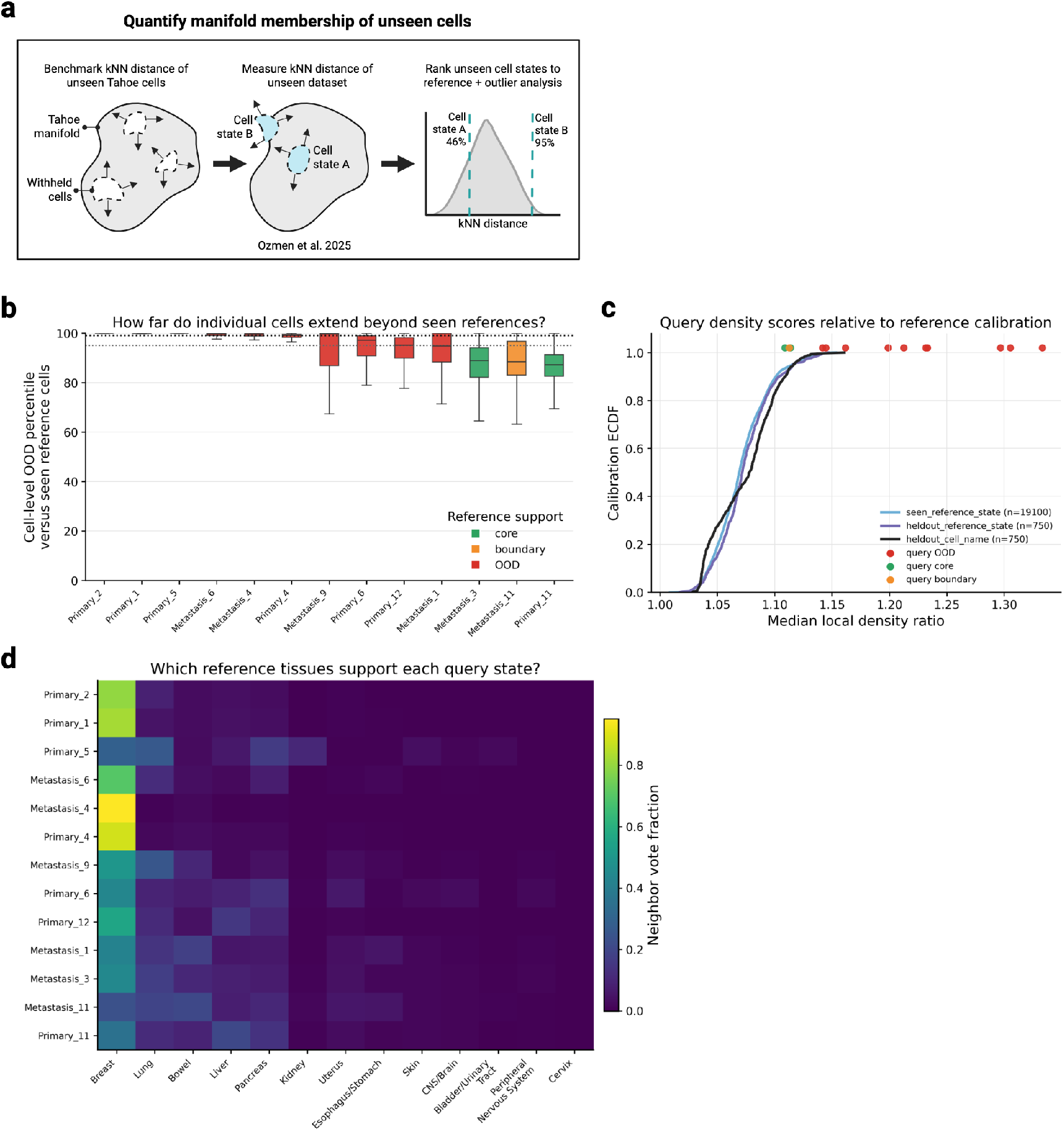
Manifold admissibility of HR+/HER2 breast cancer. **a,** Part 2 of the admissibility framework. Quantify Tahoe-100M manifold membership of unseen cell states. **b,** Cell-level manifold percentiles relative to withheld Tahoe-100M cells. Query states categorized into core (green), boundary (yellow), and out of distribution (red). **c,** Empirical cumulative distributions show median local-density ratios for seen, withheld-state, and withheld-cell-line Tahoe-100M calibrations. Larger values indicate weaker reference support. Query states are plotted above the distributions and colored by their support classification. **d,** Distance-weighted tissue annotations of the nearest Tahoe-100M reference neighbors for each query state. Rows denote query states, columns denote reference tissues and color indicates the fraction of total neighbor weight assigned to each tissue.

**Figure S10:**
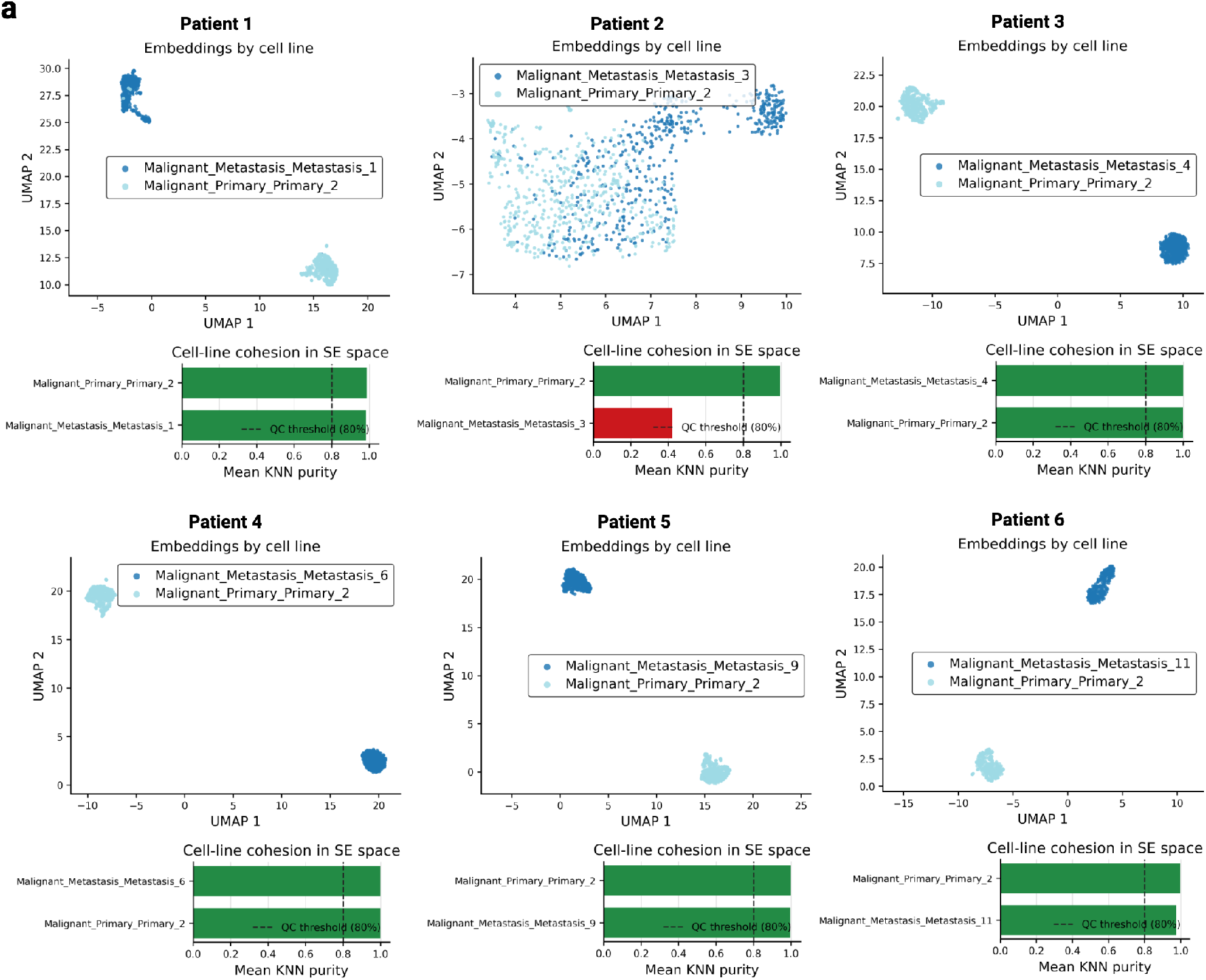
Source-target separation admissibility for all six metastatic-to-primary conversions for HR+/HER2 breast cancer. **a,** Part 3 of the admissibility framework to define a distinguishable objective for the six patients. Top: UMAP separation of source (metastatic) and target (primary) in STATE single-cell foundation model embedding space. Bottom: barplot of mean KNN purity per cell state. Dotted line indicates a purity threshold of 80%.

**Figure S11:**
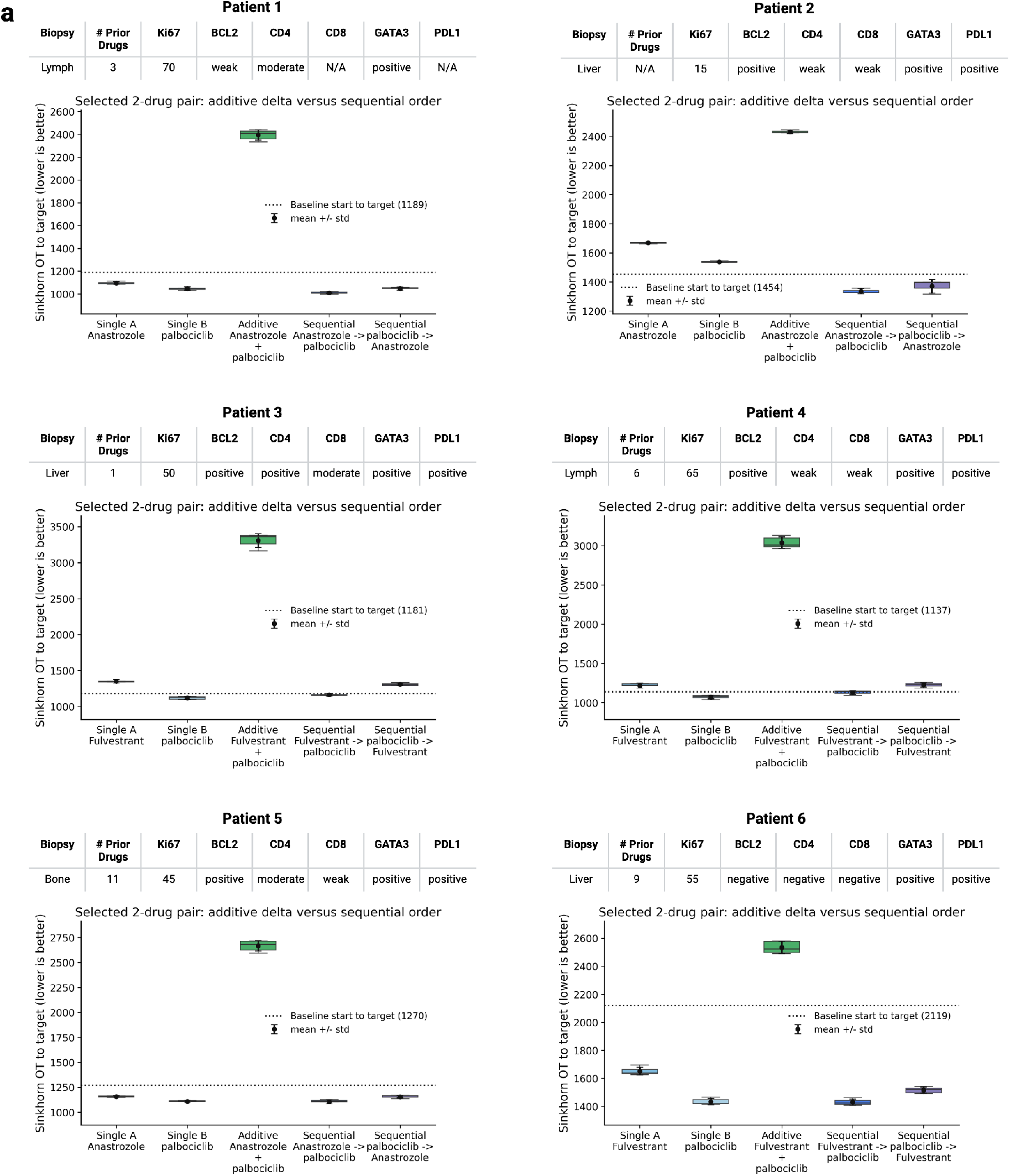
Sequential drug comparisons for all six metastatic-to-primary conversions for HR+/HER2 breast cancer. **a,** For each patient conversion from source (metastatic) to target (primary), the best 2-drug pair from the 9 FDA approved 2-drug pairs for HR+/HER2 breast cancer was selected. Boxplot of OT distance to target of starting cell state under multiple perturbation conditions of the known 2-drug combination including in order: single drug A, single drug B, additive displacement vectors of individual drug A and B, sequential A then B, sequential B then A.

**Figure S12:**
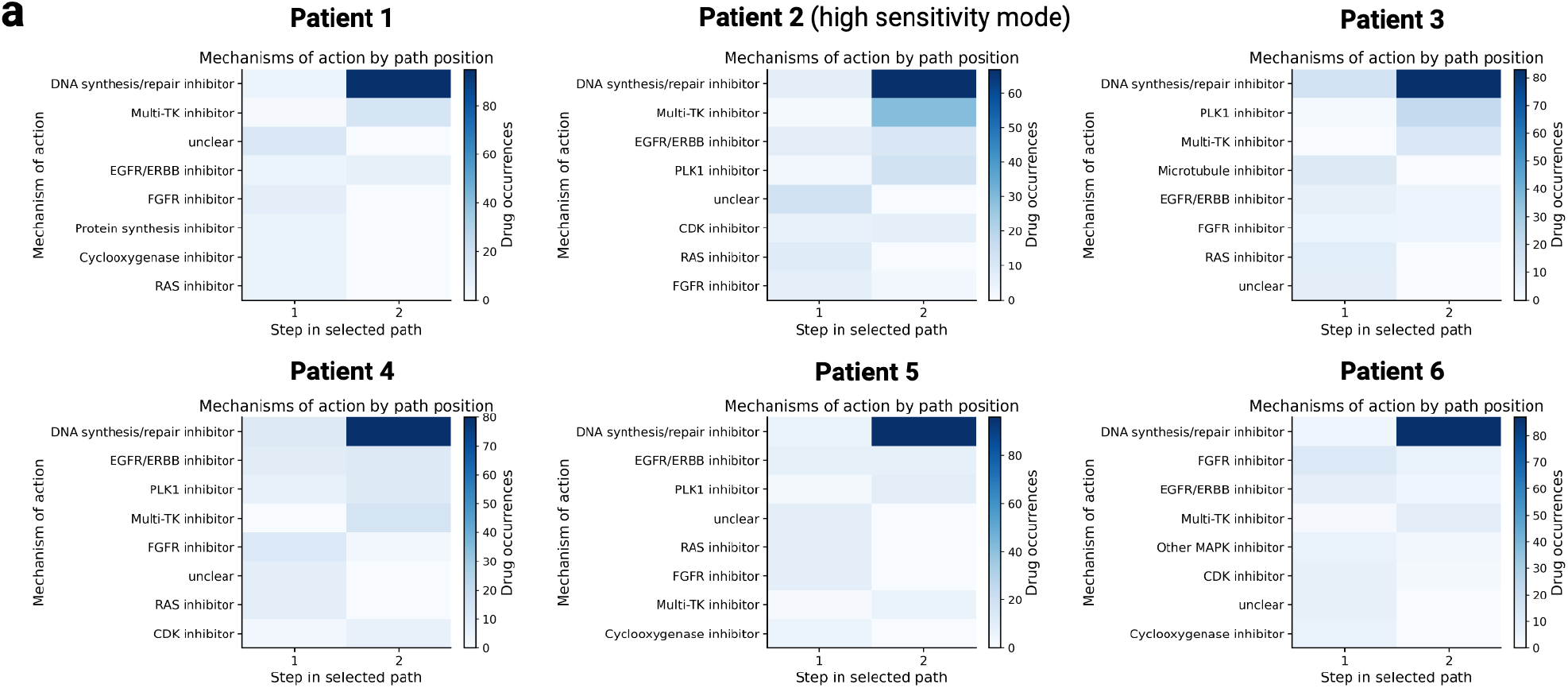
Mechanism of action frequency per patient for the conversion of metastatic malignant cells into primary tumor malignant cells. **a,** 6 patients with HR+/HER2− metastatic breast cancer. PHAROS open search conversion from metastatic malignant cells to primary malignant cells. Heatmaps, one per patient, of MOA frequencies across each step of the 2-step trajectory for the top 128 paths.

**Figure S13:**
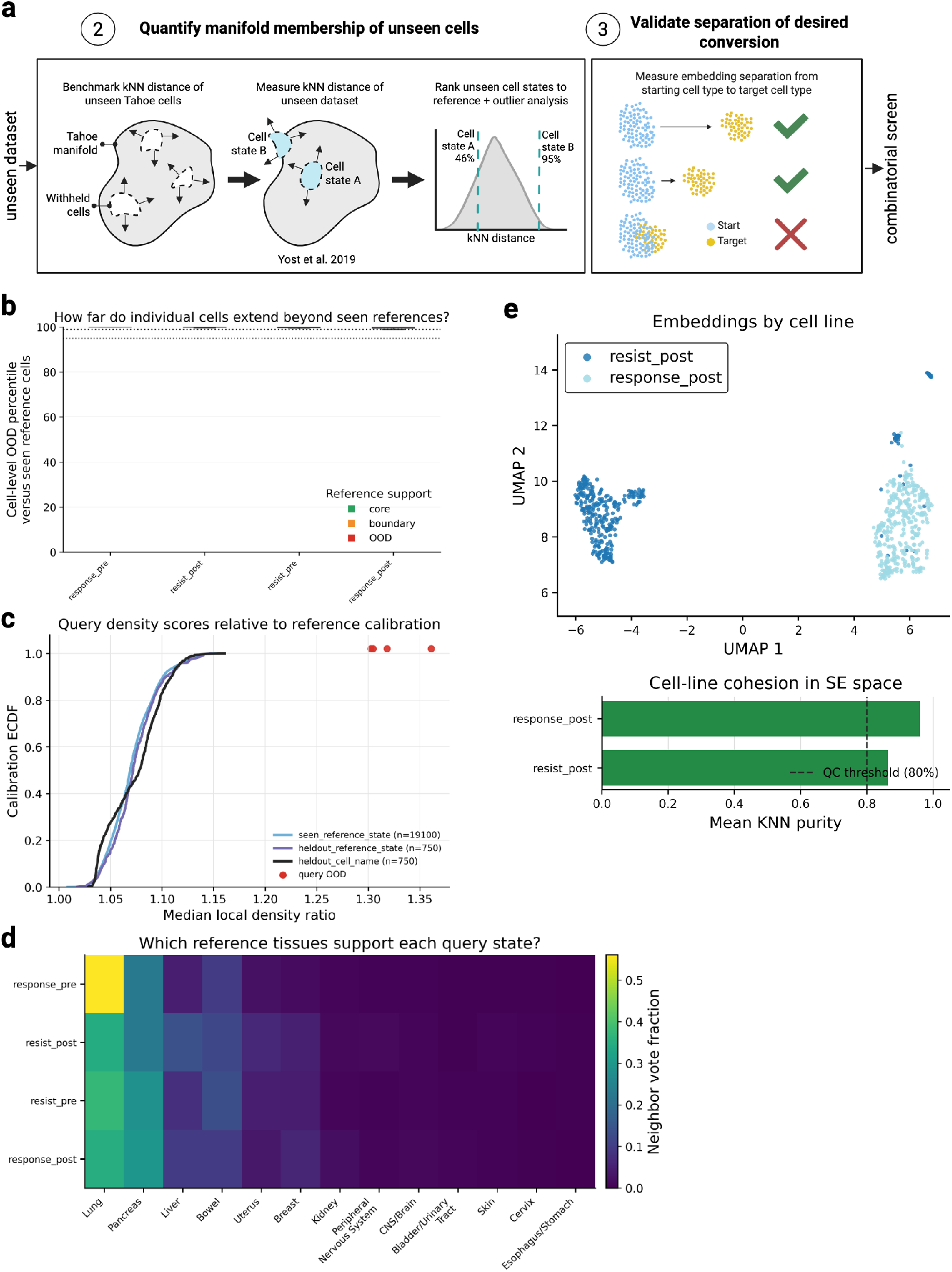
Manifold admissibility of basal cell carcinoma. **a,** Part 2 and 3 of the admissibility framework. **b,** Cell-level manifold percentiles relative to withheld Tahoe-100M cells. Query states categorized into core (green), boundary (yellow), and out of distribution (red). **c,** Empirical cumulative distributions show median local-density ratios for seen, withheld-state, and withheld-cell-line Tahoe-100M calibrations. Larger values indicate weaker reference support. Query states are plotted above the distributions and colored by their support classification. **d,** Distance-weighted tissue annotations of the nearest Tahoe-100M reference neighbors for each query state. Rows denote query states, columns denote reference tissues and color indicates the fraction of total neighbor weight assigned to each tissue. **e,** Top: UMAP separation of source (Post-immunotherapy non-responder malignant cells) and target (Post-immunotherapy responder malignant cells) in STATE single-cell foundation model embedding space. Bottom: barplot of mean KNN purity per cell state. Dotted line indicates a purity threshold of 80%.

### S2 Detailed formulation of the PHAROS conversion and search engine

This section provides an algorithmic description of the PHAROS conversion engine. Exact preprocessing, construction of the conversion-aligned PCA–PLS-DA scoring space, and experiment-specific settings are described in Methods Sections 4.5–4.9. Both hypothesis-driven evaluation and open search use the same frozen state-transition and distribution-scoring engine; they differ only in how candidate perturbation paths are generated.

#### S2.1 Cell-state conversion objective

Let **X**^(0)^ R^N×D^ denote a sampled source population and **Y** R^M×D^ a sampled target population, represented by STATE_SE_ embeddings. A perturbation label p specifies a drug and concentration from the Tahoe vocabulary. For an ordered perturbation path

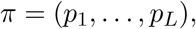

PHAROS recursively predicts the cell state after each perturbation as

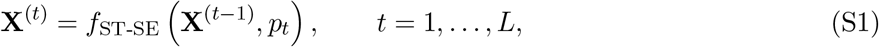

where f_ST-SE_ is the frozen pretrained state-transition model. This differs from an additive displacement approximation, in which single-drug effects are estimated from the same initial state and then summed. Consequently, PHAROS predictions may depend on perturbation order.

Let Π denote the conversion-specific scoring map. In the reported conversion-aligned analyses, Π is the fitted PCA–PLS-DA transformation described in Methods Section 4.5. Importantly, Π is used only for scoring: all ST-SE transitions in Eq. (S1) are computed in the original full SE embedding space. The terminal state produced by path π is scored against the target by entropically regularized Sinkhorn optimal transport:

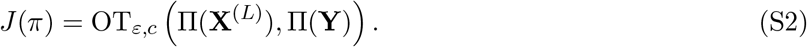

#### S2.2 Candidate generation

The hypothesis-driven and open-search modes use the same objective in Eq. (S2). In hypothesis-driven evaluation, candidate drug pairs are specified in advance. PHAROS evaluates the allowed drug concentrations and both perturbation orders and retains the configuration producing the lowest target distance.

In open search, candidate identities are not specified. PHAROS instead approximates

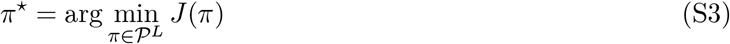

using diverse beam search, where P is the searchable set of drug–concentration perturbation labels.

#### S2.3 Robust diverse beam search

At each search depth, every retained partial path is extended by every legal next perturbation. Candidate states are first scored using a 10-iteration Sinkhorn approximation. The best αB candidates are then recomputed with the full 100-iteration Sinkhorn calculation.

To reduce sensitivity to the particular cells sampled in the source and target populations, candidates surviving this prefilter are replayed across multiple source–target batches. For a candidate path π, let J_b_(π) denote its Sinkhorn target distance in batch b. PHAROS uses the robust score

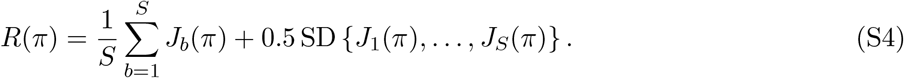

A standard beam search can become dominated by highly similar paths where drugs used are the same with varying concentrations. For candidate π and the set _sel_ of paths already selected into the new beam, PHAROS defines

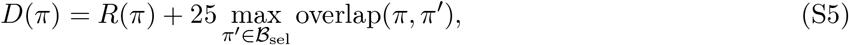

where overlap(π, π^′^) is the fractional overlap in base-drug identities between the two paths. When _sel_ is empty, the overlap penalty is zero. Candidates are added sequentially according to increasing D(π) until the beam is full.

For all reported two-drug open searches, the beam width was B = 128, the maximum depth was L_max_ = 2, and the prefilter multiplier was α = 10.

#### S2.4 Path constraints

Perturbation labels corresponding to DMSO were excluded from the searchable library. Different concentrations of the same drug are distinct perturbation labels, but once a base drug appears in a path, no other concentration of that drug may be selected later in the same path.

##### Algorithm 1 PHAROS robust diverse beam search

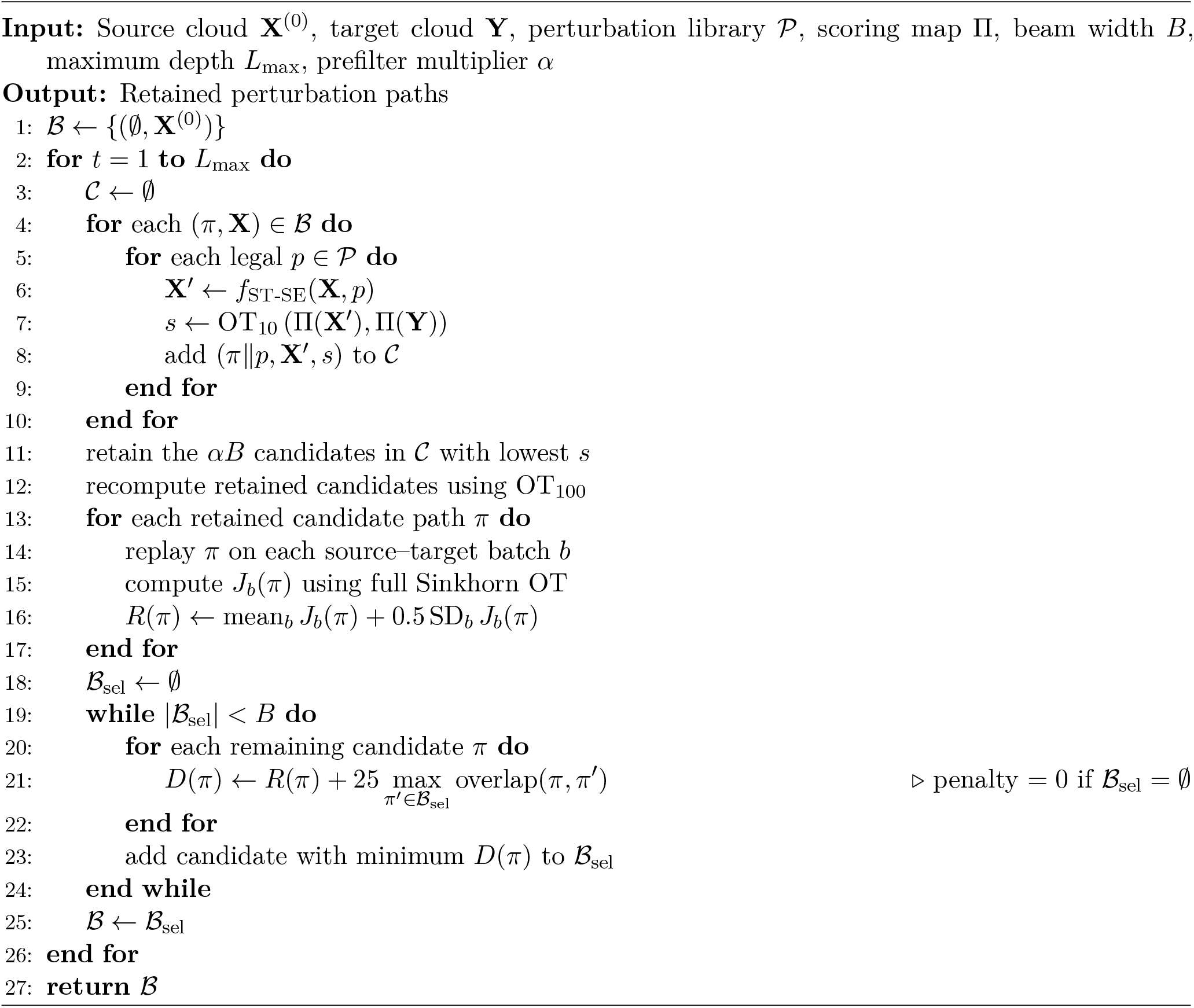

Here π p denotes appending perturbation p to path π. Drug–concentration labels are treated as distinct actions during expansion, whereas path-reuse constraints and the diversity penalty operate on base-drug identity.

## S3 Model validation - interpreting the model given our framework

**Figure S14:**
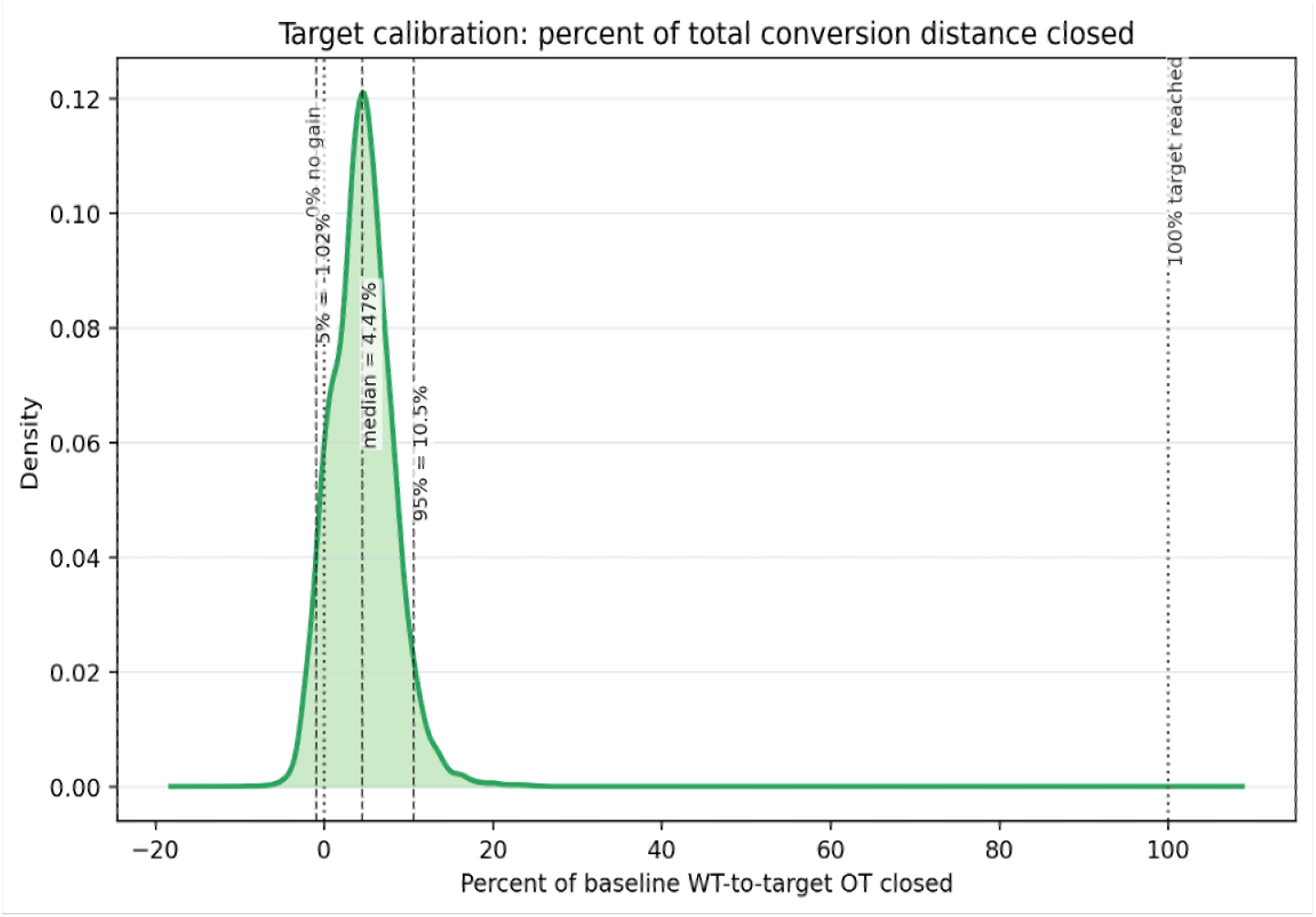
Target calibration in full-d space. Density plot of the percentage of the initial WT-to-target optimal-transport (OT) distance (in full-d space) closed when a group of WT cells is passed through the model with the ground-truth drug as the perturbation label. Each observation corresponds to a combination of drug and cell-line across Tahoe, with the target being the true perturbed population for that combination. Read S4 to understand better the difference in results between the compressed and the full-d space.

**Figure S15:**
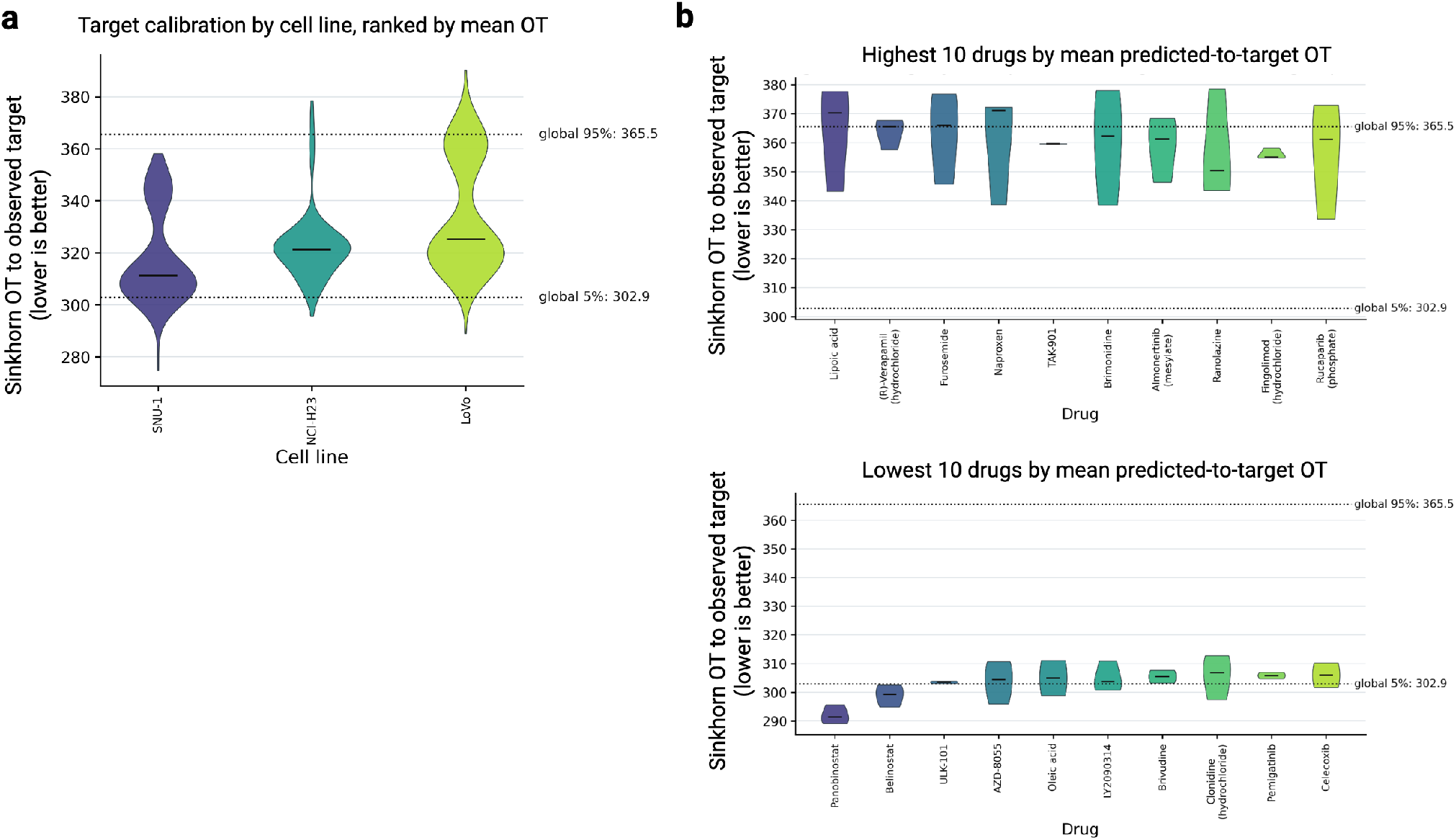
Target calibration by cell line and drug. **a,** Density plots of OT distance between a group of WT cells passed through the model with the ground-truth drug as the perturbation label and a group of experimental cells perturbed with the same drug. Each observation corresponds to a combination of drug and cell-line across Tahoe, and the data is grouped by cell-line. **b,** Same as in (a) but grouped by drug and showing only highest and lowest drugs by mean OT distance predicted-to-target.

**Figure S16:**
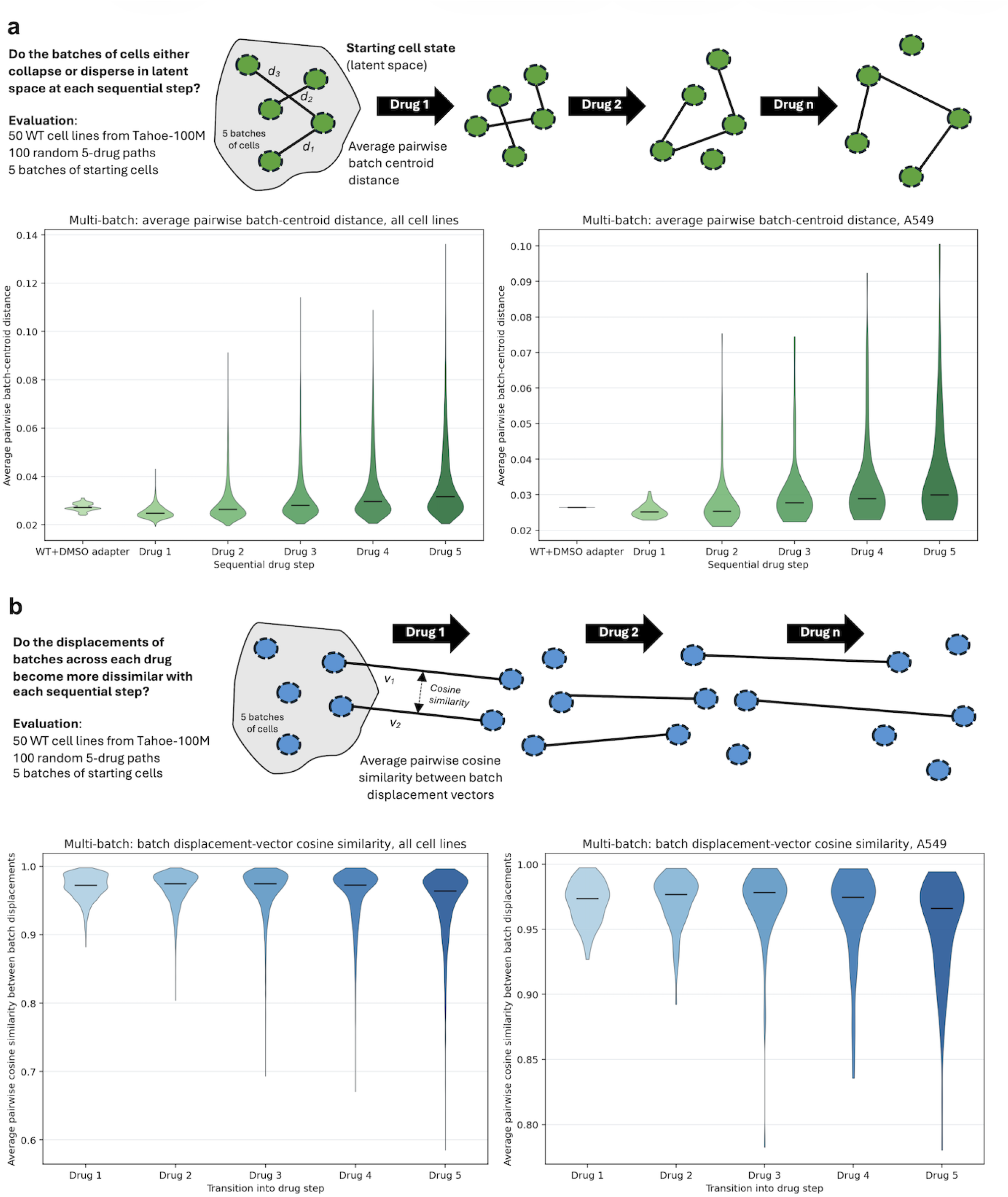
Stability of sequential ST-SE rollouts across drug steps. **a-b,** Evaluated over all 50 WT (no drug) cell lines from Tahoe-100M across 5 batches of starting cell states and 100 random 5 drug step trajectories. **a,** Quantifying how the distribution of batches evolves at every step. Distribution of average pairwise batch-centroid distance at each step of the 5 drug trajectories. Left: all cell lines. Right: A549. **b,** Quantifying how similar the batches are displaced with each sequential drug. Distribution of average pairwise cosine similarity between batch displacements comparing cell states in drug N and drug N-1. Left: all cell lines. Right: A549.

**Figure S17:**
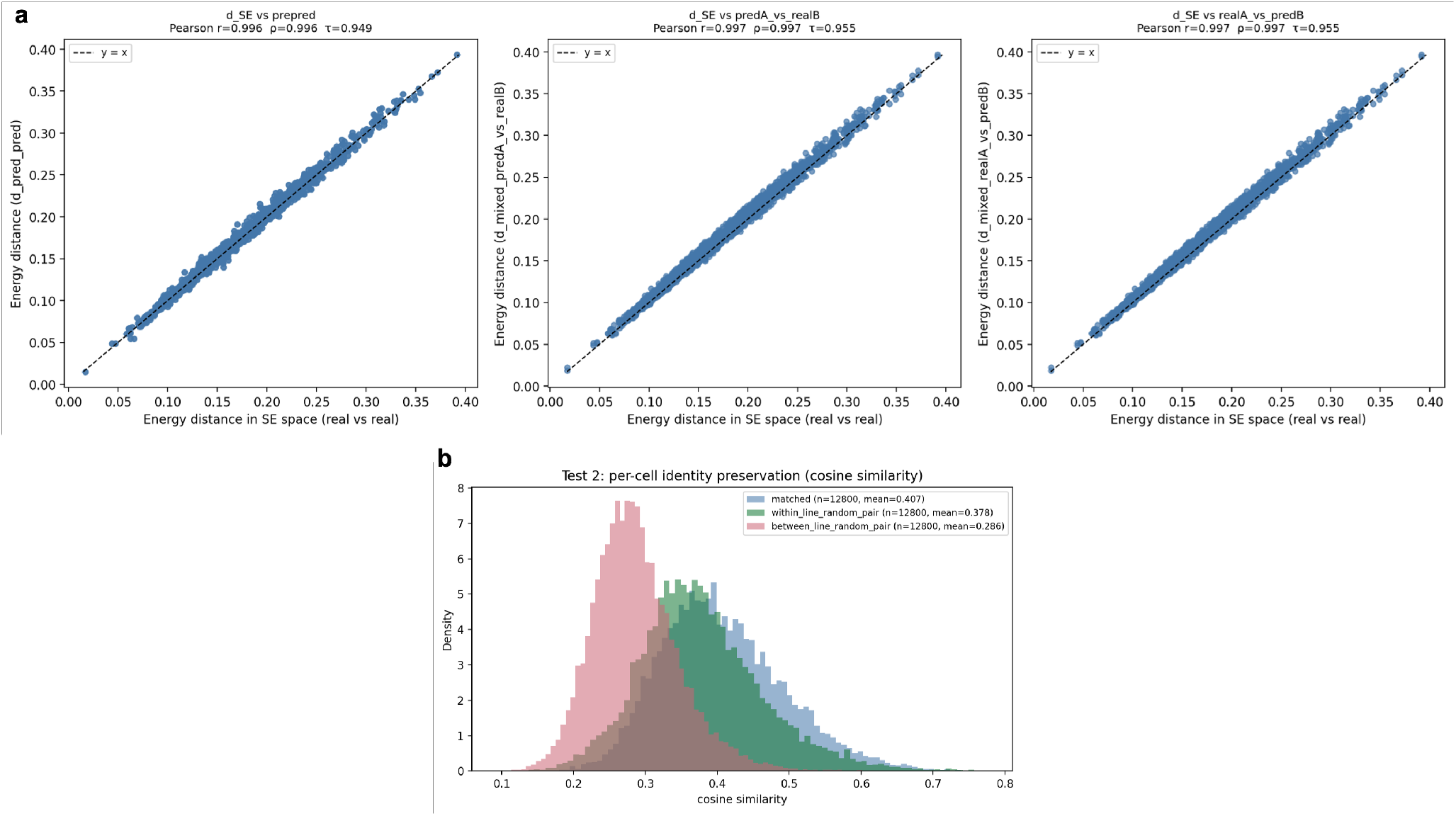
Alignment for unperturbed cells between SE and ST-SE. **a,** The first scatter plot shows the correlation between the energy distance of two groups of 256 unperturbed cells from different cell lines computed in SE space and the same metric in ST-SE space, that is after the DMSO adapter. In the other two scatter plots only one of the two groups is passed through the ST-SE model (DMSO adapter), while the other is kept in SE space. **b,** The overlaid density histogram compare the distribution of cosine similarity across three groups. In the matched group, for each cell we compute the cosine similarity between its unperturbed SE embedding and the ST-SE output for the same cell after passing it through the predictor with DMSO as the perturbation label. In the within-line random pair group, we compute cosine similarity between SE embeddings of two different unperturbed cells sampled from the same cell line (neither passed through ST-SE). In the between-line random pair group, we compute cosine similarity between the SE embedding of a cell in one line and the SE embedding of a cell drawn from a different cell line.

## S4 Understanding the effect of dimensionality reduction

In this section, an informal noise model is presented, which serves to motivate the use of dimensionality reduction techniques, such as PCA– PLS-DA, to enhance the match between the predicted and target cell populations. This provides a theoretical explanation for the differences observed between the compressed and full-dimensional spaces. The Sinkhorn optimal-transport distance metric is employed for the analysis, with a primary focus on the cost matrix.

Consider two empirical cell populations

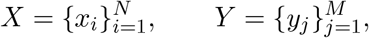

where each cell embedding is decomposed into a biologically informative component and a nuisance component:

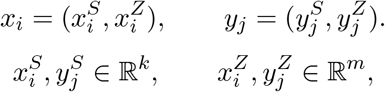

with m = D k. We assume that the nuisance coordinates have the same distribution in the two populations,

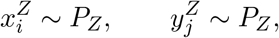

thus do not contain useful information for distinguishing the two states.

Using the squared Euclidean cost, the pairwise cost between cells x_i_ and y_j_ is

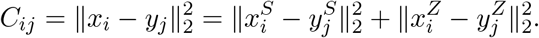

We define

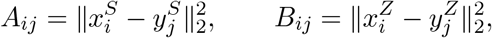

so that

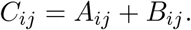

A_ij_ is the biologically relevant contribution to the cost, whereas B_ij_ is induced by nuisance variation.

Assume that the nuisance coordinates are independent and have equal variance σ^2^. For a single nuisance coordinate, define

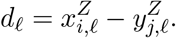

Since and *x^Z^_j.l_* are independent and identically distributed,

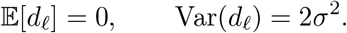

Therefore,

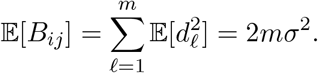

If the nuisance coordinates are Gaussian, then

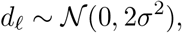

and hence

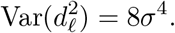

It follows that

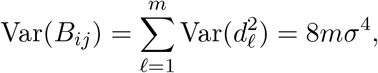

and

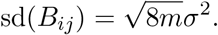

Thus the nuisance contribution can be decomposed as

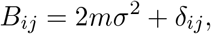

where

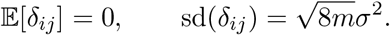

While the constant term 2mσ^2^ shifts all pairwise costs by the same amount, the problematic component is the fluctuation δ_ij_, because it can change the relative ordering of the entries of the cost matrix.

Informally, for population matching to be driven by biological structure rather than nuisance variation, differences in signal costs should dominate nuisance fluctuations. For two candidate cell pairs (i, j) and (i^′^, j^′^), this requires

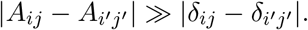

Since the scale of δ_ij_ is

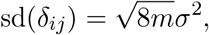

a useful informal signal-to-noise ratio is

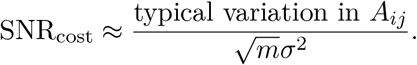

As the number of nuisance dimensions m increases, this ratio decreases, and a low SNR can make pairwise costs noisy and distort population matching.

Since PCA–PLS-DA reduces dimensionality by selecting directions that best discriminate between the two populations, it can significantly reduce nuisance variation while preserving signal associated with the population-level contrast. This motivates computing Sinkhorn OT scores after applying a supervised dimensionality reduction step, especially when embeddings contain lots of dimensions that are weakly informative for the conversion of interest.

## S5 CPA vs STATE

We also considered using CPA within PHAROS as the perturbation predictor, primarily due to the model structure, which, at least theoretically, allows drugs to be applied simultaneously in latent space, thereby eliminating the sequential application strategy required by our ST-SE implementation. Furthermore, to our knowledge, CPA is the only model that can be effectively applied to predict multiple drug perturbations with single-cell resolution. However, we encountered significant challenges in identifying a solution to two substantial limitations. Firstly, CPA lacks the capacity to regulate dataset-specific effects, a capacity that is partially addressed by STATE via the utilization of the DS token in the SE module. Secondly, in its standard formulation, CPA does not possess the capability to predict the effects of perturbations in unseen cell lines because cell-line identity is encoded via categorical covariate embeddings that are undefined at inference for unseen lines. To solve such limitations one could modify the architecture to eliminate both cell line embeddings and the adversarial task of predicting cell lines, and train using SE embeddings both for the model’s input and output spaces. However, this would significantly deviate from the original CPA architecture and demand large-scale retraining on the whole Tahoe corpus, whereas PHAROS relies on a pretrained ST-SE model.

## S6 Interpretation of partial positive-control recovery

In some positive controls, the experimentally applied drug pair does not lie in the lower tail of the OT-distance-to-target distribution of random two-drug perturbations. Several non-exclusive explanations may account for this. First, ST-SE is far from a perfect perturbation predictor. Error compounds as we predict sequentially, especially when at the intermediate step the cell moves to an out-of-distribution part of the space. Furthermore PHAROS applies drugs sequentially (A → B), whereas positive controls are based on simultaneous co-treatment (A + B). In certain scenarios order may matter, that is A → B (and/or B → A) is substantially different from A + B. For example, this might be the case if two drugs inhibit parallel, partially redundant branches of a regulatory network. Lastly, PHAROS might struggle even more when dealing with drug combinations characterized by synergistic or antagonistic behaviors, that is combinations whose joint effect deviates from the addition of single-drug responses. Importantly, at least in theory, a sequential predictive approach might help in this case as the second drug is applied conditional on the cell cloud produced by the first. This advantage assumes that the intermediate state is predicted accurately and remains in-distribution for ST-SE.

## S7 Proposed reinforcement-learning search (not evaluated)

Beam search was used for all experiments reported in this study. The major disadvantage is that such an algorithm is greedy with respect to the distance metric at each step and can miss paths requiring temporarily suboptimal intermediate perturbations that position the cloud embeddings in a favorable part of the space for later moves. To tackle this problem, drug-path discovery can be formulated as a finite-horizon Markov decision process (MDP) in which the pretrained ST-SE model defines the transition dynamics. We outline this formulation for completeness, but it was not implemented.

### S7.1 MDP formulation

We model sequential drug selection as an MDP (S, A, P, R, γ).

**Environment state.** At step t, the environment maintains a cell cloud **X**_t_ R^N×D^. A fixed target cloud **Y** R^M×D^ is sampled once per episode from a target state. The starting cloud **X**_0_ is sampled from the starting state.

**Actions.** An action a_t_is a perturbation label (drug and concentration) from the ST-SE vocabulary. The same path constraints as in Section S2.4 apply.

**Transitions.** Transitions are determined by the sequential application of ST-SE:

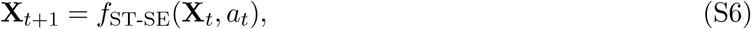

as in Eq. (**??**). Given **X**_t_ and a_t_, the next state is deterministic. Episodes have maximum length L_max_.

**Reward.** We define a reward function using Sinkhorn optimal-transport (OT) distance to the target cloud **Y**:

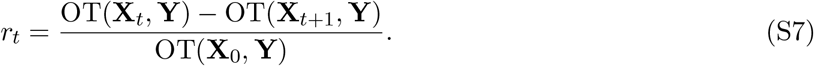

Positive reward therefore corresponds to progress toward the target distribution.

### S7.2 Goal-conditioned Q-network

We propose a single action-value network Q(**s**_t_, a; ***θ***) trained across many start–target pairs. The model is goal-conditioned, that is the target enters the state, allowing for inference on any start–target couple of interest.

**State encoding.** Because the Q-network requires a fixed-size input, the full cell cloud **X**_t_ is summarized by a map ψ(·) applied to the current and target clouds. For example,

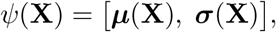

where ***µ*** and ***σ*** are the mean and standard deviation (square roots of the diagonal entries of the empirical covariance matrix) across cells. Optionally, ψ can be computed after a fixed global dimensionality reduction of SE embeddings (e.g. PCA fit on all Tahoe cells embedded with SE).

Let **h**_t_ = ψ(**X**_t_) and **g** = ψ(**Y**) (computed once per episode). The input to the Q-network at step t is

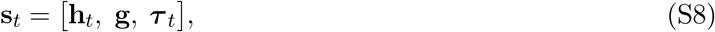

where ***τ*** _t_ = **e**_Lmax_−t ∈ {0, 1}^Lmax+1^, **e**_k_ is the kth standard basis vector.

**Learning.** Following the deep Q-network (DQN) framework, we approximate Q^∗^(**s**, a) with a neural network Q(**s**_t_, a; ***θ***) optimized by temporal-difference learning. Illegal actions are masked when computing max_a_*′* Q(**s**_t+1_, a^′^; ***θ***^−^). A periodically updated target network ***θ***^−^ stabilizes training. As with standard function approximation, convergence to Q^∗^ is not guaranteed.

**Inference.** At inference, given a starting and a target state of interest, the trained action-value network Q(**s**_t_, a; ***θ***) constructs a predicted path by choosing a_t_= arg max_a∈Alegal(_**_s_***_t_*_)_ Q(**s**_t_, a; ***θ***) at each step and updating **X**_t+1_ = f_ST-SE_(**X**_t_, a_t_). Alternatively, the trained neural network can be employed within a tree-search procedure (e.g. beam search) to search for additional promising paths.

### S7.3 Training data (proposed)

Learning requires a diverse set of simulated experiences sampled within the Tahoe manifold, otherwise ST-SE would have to predict out-of-distribution transitions. In practice, one would select a collection of start–target cell-line pairs whose SE embeddings span this space in a representative way. Episodes would then be generated by rolling out ST-SE from sampled starting clouds toward fixed target clouds, with OT-based rewards as in Eq. (S7), and actions chosen ɛ-greedily: with probability ɛ a legal perturbation is sampled uniformly at random, and with probability 1 ɛ the greedy action arg max_a∈Alegal(_**_s_** _)_ Q(**s**_t_, a; ***θ***) is chosen. We do not specify a particular pair-selection, but the main requirement is coverage and diversity of (c_s_, c_t_) tasks within the Tahoe manifold.

### S7.4 Relation to beam search

Both approaches use the same ST-SE transition model and OT-based notion of progress toward a target distribution. Beam search evaluates explicit multi-step paths without learning and was used for all results in this study. Reinforcement learning could in principle yield a reusable policy across conversion tasks, but each environment step requires a full ST-SE forward pass on N cells, and training would require many simulated episodes. We therefore leave this as a direction for future work.

